# Taxon-resolved sources of organic carbon preserved in lake sediments using the sedaDNA GenC pipeline

**DOI:** 10.64898/2026.05.21.726723

**Authors:** Zijuan Yong, Josefine Friederike Weiß, Kathleen R. Stoof-Leichsenring, Sisi Liu, Ulrike Herzschuh

## Abstract

Organic carbon (OC) burial in lakes is an important component of the global carbon cycle, but the source organisms of preserved OC remain poorly resolved. Here we develop the genC pipeline, which combines sedimentary ancient DNA concentrations, read-based taxonomic assignments, and group-specific priors for DNA and cellular carbon content to derive OC_DNA-projected_, a semi-quantitative proxy for the magnitude and taxonomic composition of preserved sedimentary OC. We apply genC to six high-latitude lake records spanning the last 30,000 years. OC_DNA-projected_ broadly agrees with independent proxies for total organic carbon and aquatic contribution, supporting its reliability. Our results indicate that environmental conditions, especially warming, rather than preservation alone, are the main drivers of preserved OC variation. Terrestrial sources, mainly woody plants, dominate lake sediment OC. Eukaryotic algae as well as aquatic and terrestrial bacteria become more important during the warmer Holocene. These results establish sedaDNA as a taxonomically resolved tool for reconstructing long-term changes in preserved lake-sediment OC.

## Introduction

Although lakes cover only a small fraction of the Earth’s surface, they bury organic carbon (OC) at disproportionately high rates per unit area^1^ and therefore play an important role in regional and global carbon cycling^2, 3^. Burial transfers organic matter from the water column and catchment to sediments, where a fraction is retained on centennial to millennial timescales^4, 5^. Lake sediments thus archive both depositional inputs and post-depositional modification including degradation^6^, but the taxonomic composition of the preserved OC pool remains difficult to resolve over long timescales. Recent increases in lake OC burial^7^, including in northern regions, further emphasize the relevance of lacustrine sediments for long-term carbon storage.

Climate influences lake carbon burial by, among other factors, affecting in-lake productivity, catchment runoff and land cover, and sediment delivery^3^. Higher OC burial has, for example, been observed during warm Holocene intervals in northern high-latitude lakes^7^. At the same time, mineral availability^5^, sedimentation rate^8^, and taxonomic composition influence organic matter preservation in sediments. Disentangling depositional from preservational effects on burial therefore remains challenging, particularly when the aim is to separate these influences for the aquatic and terrestrial fractions of the preserved sedimentary OC pool.

A broad toolkit has been used to investigate the organismic sources of the preserved sedimentary OC pool, including bulk elemental and isotopic proxies^9^, such as carbon-to-nitrogen (C/N) ratios and δ¹³C values^10^; biomarkers and compound-specific isotope analyses^11^; HPLC-based algal pigments^12^; FTIR spectroscopy ^13^; hyperspectral methods^14^; and pyrolysis GC–MS^15^. For instance, lignin biomarkers can differentiate and quantify the relative contributions of gymnosperm versus angiosperm plant sources^16^. Similarly, isotope signals from fatty acids are useful for assessing aquatic plant contributions and their productivity^17^. Additionally, glycerol dialkyl glycerol tetraethers (GDGTs) serve as microbial biomarkers, and branched GDGTs (brGDGTs) can indicate contributions from specific microbial sources under certain environmental conditions^11^. The ratio of steranes to hopanes is frequently applied to evaluate the relative inputs of eukaryotic versus bacterial biomass to sedimentary archives^18^, while phospholipid fatty acid (PLFA) analysis helps quantify viable microbial biomass^19^. These approaches have yielded important insights into OC dynamics in lake sediments, but most remain limited in their ability to resolve the relative contributions of many organism groups simultaneously and at relevant taxonomic resolution. There is therefore substantial value in developing complementary taxon-resolved approaches that can be used alongside established biogeochemical proxies.

Sedimentary ancient DNA (sedaDNA) analysis has greatly advanced ecosystem reconstructions at high taxonomic resolution. Sedimentary metagenomics, involving the extraction and shotgun sequencing of total DNA from sediment, offers the potential to detect a broad range of taxa represented by DNA in a sample^20^. Unlike metabarcoding, which depends on PCR amplification of specific marker genes, shotgun metagenomics employs an untargeted approach, facilitating more comprehensive assessment of multiple taxonomic groups from a single dataset^21^. For instance, this method enabled integrated reconstructions of terrestrial and aquatic ecosystem responses to glacial–interglacial climate transitions^22^. Additionally, von Hippel et al.^23^ utilized sedimentary metagenomics to reconstruct postglacial shifts in plant, fungal, and bacterial communities linked to soil development and vegetation change. However, sedimentary ancient DNA analyses have rarely been applied explicitly to trace organism-specific sources of preserved sedimentary organic matter. Zimmermann et al.^24^ evaluated plant-derived organic matter composition in permafrost sediments, while Herzschuh et al.^20^ traced terrestrial plant-derived organic contributions to marine sediments using terrestrial ancient DNA as a proxy for terrigenous organic matter burial. However, variation in DNA extraction efficiency among taxa^25^ and differences in sequencing read recovery^26^ can introduce substantial bias into metagenomic approaches.

Beyond compositional assessments, recent studies have used eDNA to derive quantitative estimates of organismal abundance or biomass, for example in fish communities^27^. Likewise, sedimentary metagenomics holds promise for first-order, taxonomically resolved estimation of the preserved sedimentary OC pool. However, sedimentary DNA preservation remains incompletely understood^21^. DNA can persist through adsorption to mineral surfaces^28^, within shielded cells^6^, or in other stabilized pools. It is still poorly understood to what extent, and over which timescales, DNA preservation tracks bulk organic matter decay or shares similar first-order controls (e.g., ^21, 28, 29, 30^). These factors, together with taxon-specific variation in DNA extraction efficiency and sequencing read recovery, can skew estimates of the contribution of taxa to total organic carbon. In addition, robust semi-quantitative reconstructions require the integration of taxon-specific DNA content per cell^31^ and organism-specific carbon mass per cell^32^. Thus, there is a need for a framework capable of combining these factors to convert DNA concentration into semi-quantitative carbon-content estimates, here termed OC_DNA-projected_.

Here, we develop and apply the genC pipeline (Fig. 1) to sedimentary ancient DNA metagenomic data from six high-latitude lake sediment cores from Siberia and Alaska spanning at the last 30,000 years (Fig. 2a) (MIS3 to MIS 1). By combining sedimentary DNA concentrations, read-based taxonomic assignments, and group-specific priors for DNA content and cellular carbon content, we derive OC_DNA-projected_ as a sedaDNA-based estimate of taxon-specific contributions to preserved sedimentary OC. We compare OC_DNA-projected_ with established geochemical and biological indicators, examine variations in relative aquatic and terrestrial contributions, and use the data to describe temporal variation in the taxonomic composition of the preserved OC pool. We interpret genC as a first-order, semi-quantitative, taxonomically resolved framework for reconstructing the composition of preserved sedimentary organic carbon and thus provide a novel opportunity for exploring past pre-and postdepositional processes governing organismal sources of carbon burial.

**Figure 1.**
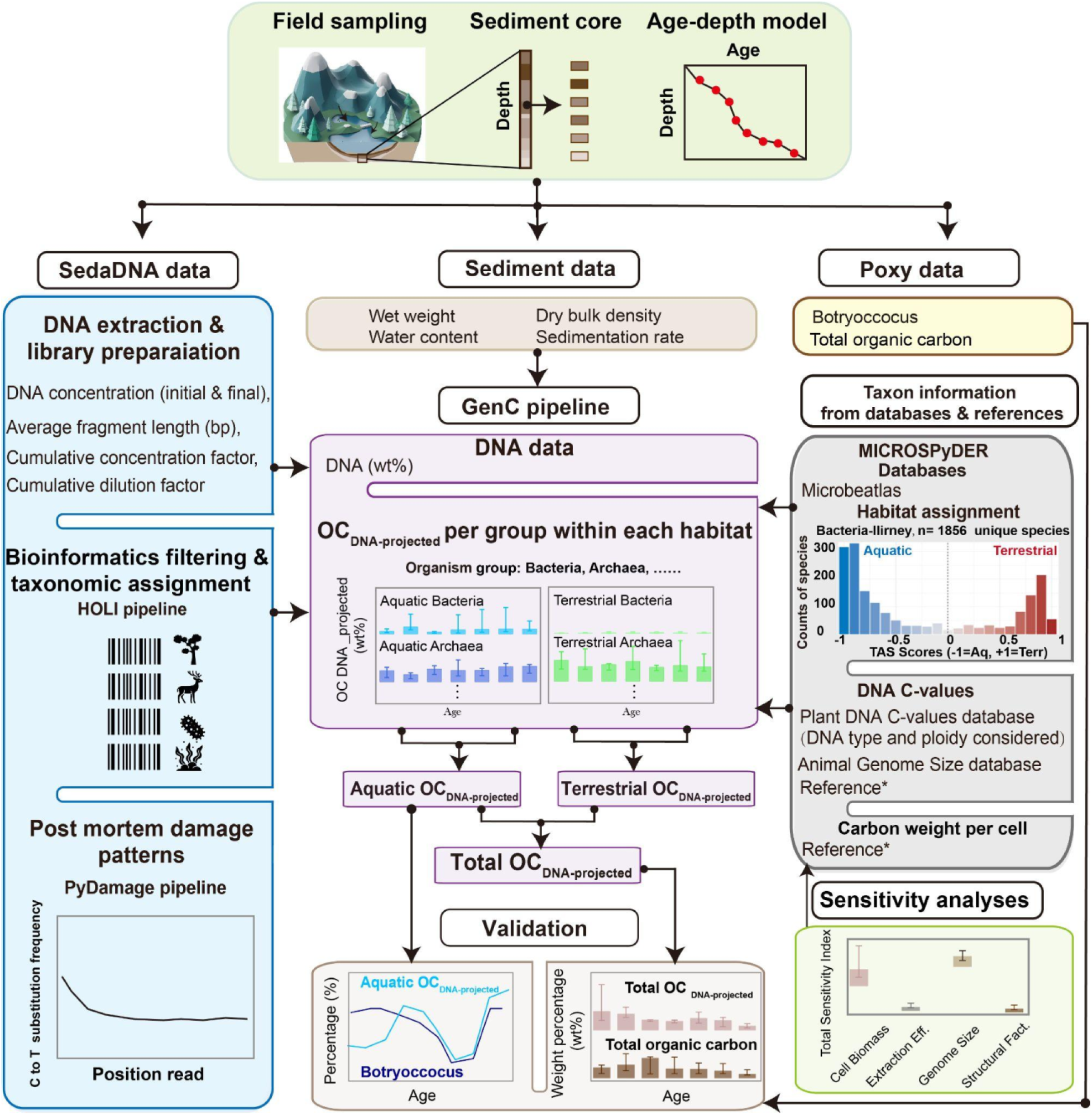
General framework illustrating sedaDNA-projected organic carbon estimation and validation (this study). The flowchart outlines the workflow for quantifying the main contributors of lake organic carbon based on sedimentary ancient DNA (sedaDNA). The workflow includes field sampling, establishing an age-depth model, DNA extraction and library preparation, followed by bioinformatic filtering and taxonomic assignment. Post-mortem damage patterns are also used to evaluate the quality of sedimentary DNA. Library preparation data combined with sediment data to calculate DNA data. After taxonomic assignment, species-level habitat classification is carried out using reference databases. To estimate DNA-projected organic carbon, we combine the DNA weight percentage of organism groups with DNA C-values and DNA data. The resulting sedaDNA-based organic carbon estimates are validated against geochemical proxies with total organic carbon (TOC) measurements. *Reference sees (Supplementary Data 4).

**Figure 2.**
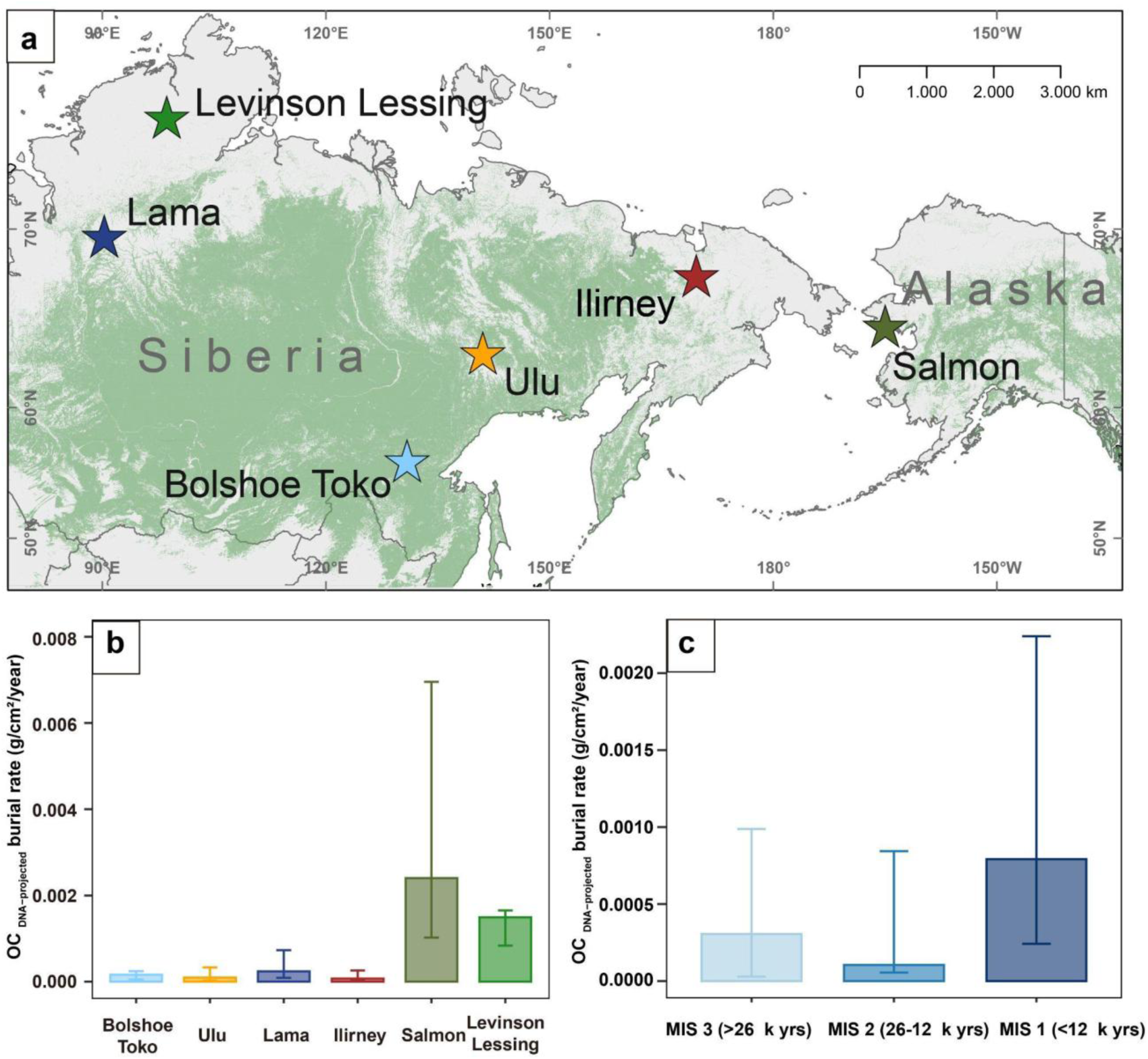
Information about location of the study sites and site-specific estimates of organic carbon DNA projected burial rate. **(a)** Map showing the locations of the sediment cores from Lake Lama, Lake Levinson Lessing, Lake Bolshoe Toko, Lake Ulu, Lake Ilirney, and Lake Salmon. **(b)** Organic carbon DNA-projected burial rate (g/cm^2^/year) for every lake. **(c)** Median Organic carbon DNA-projected burial rate (g/cm^2^/year) across all lakes during Marine Isotope Stages (MIS 1–3). For panels **(a)**, green shading indicates the current distribution of evergreen needle-leaved boreal forests.; For panels **(b, c)**, the top of each bar represents the median of samples, while the lower and upper ends of the error bars indicate the first (Q1) and third (Q3) quartiles of those samples.

## Results and discussion

### Sedimentary metagenomic data quality from 6 lake sediment records

We used published palaeogenomic data from 6 Lake sediment cores generated from shotgun sequencing^33^. In total, 16,785,147 reads were obtained from Lake Ilirney, 17,607,944 from Salmon Lake, 78,593,960 from Lake Levinson-Lessing, 93,565,199 from Lake Lama, 54,335,935 from Lake Bolshoe Toko, and 11,941,141 from Lake Ulu.

Of the quality-filtered reads, 89–94% could be taxonomically assigned at least at the superkingdom, 71–77% the family level and 18% to 31% at the species level (Supplementary Data 1). Reads were categorized into major taxonomic groups with Bacteria (40.65%), Viridiplantae (47.60%, further separated into woody, non-woody, and aquatic plants), Archaea (1.88%), Metazoa (0.86%), aquatic algae/protists (0.68%), Fungi (0.10%), Viruses (0.13%).

The wet-lab results from the blank controls confirmed very clean sample processing and no evidence of substantial contamination from laboratory reagents. All extraction blanks (controls for extraction chemicals) showed no or only very low DNA concentrations (0–0.17 ng µL⁻¹). A few library blanks (controls for library preparation chemicals) showed measurable DNA concentrations after indexing PCR, likely due to amplification of library adapters rather than sample DNA (Supplementary Data 2). Detected sequence reads across all blanks were very low, accounting for only 0.16% of the total read counts across samples and blanks. For Bacteria, the median proportion of read counts in the blanks was 0.033%, compared to 0.014% for Eukaryota, 0.00022% for Archaea, and 0.000041% for Viruses.

The ancient origin of DNA reads recovered from the samples was supported by characteristic post-mortem damage patterns. Filtered contigs from the seven taxonomic groups in each record show the typical increase in C-to-T substitutions toward the 5′ ends (Supplementary Fig. 1), consistent with the ancient origin of the DNA fragments^34^. Across all taxonomic groups and lake sites, we found a mean proportion of 54.9% damaged reads associated with the filtered contigs (Supplementary Fig. 2), which is broadly similar to previous sedimentary ancient DNA studies^23, 35^. Median read-damage proportions were highest for Archaea (74%), Bacteria (60%), Metazoa (59%), and Viridiplantae (55%), whereas slightly lower values were detected for aquatic algae/protists (52%), Viruses (52%), and Fungi (43%). The relationship between C-to-T substitution frequency at the first position and sediment age was examined exemplarily for Lake Lama and supports increasing damage with sediment age for most taxonomic groups, except fungi (Supplementary Fig. 3, see also von Hippel et al.^23^).

### Reconstruction of sedimentary DNA-projected organic carbon (OC_DNA-projected_) using GenC pipeline

We calculated the DNA weight percentage (wt%) from measured DNA concentrations relative to the dry sediment mass (following Herzschuh et al.^20^, Fig. 1). The resulting DNA wt% ranged from 0.0000017 to 0.035 wt%, with a median of 0.001 wt% (Supplementary Data 3), and show a clear increase from the glacial period toward the Holocene (Fig. 3a).

**Figure 3.**
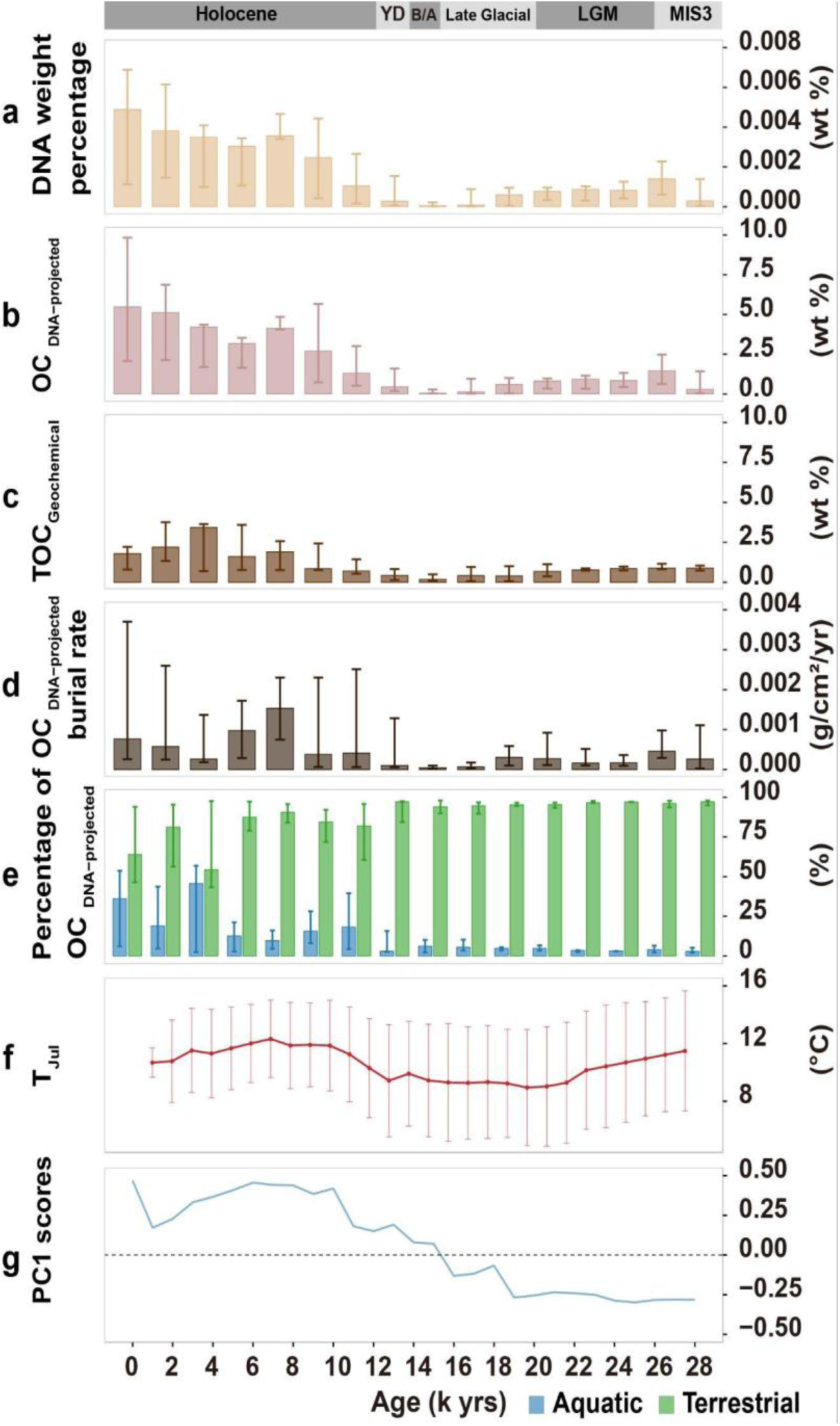
DNA-projected, biogeochemical organic carbon from six lake sediment cores during the last 30,000 years and potential drivers. **(a)** DNA weight percentage (wt%) change over time; **(b)** OC_DNA-projected_ weight percentage (wt%) change over time; **(c)** geochemical data of total organic carbon (TOC) weight percentage (wt%) change over time; **(d)** OC_DNA-projected_ burial rate (g/cm²/years) change over time; **(e)** terrestrial and aquatic OC_DNA-projected_ weight percentage (wt%) change over time; **(f**) reconstructed mean July temperature (°C) based on six pollen sites in the study region using the weighted averaging partial least squares (WAPLS) method, over the past 30,000 years, Points and line show median values across records, and error bars represent ±1 standard deviation (SD); **(g)** the first axis scores of a principal component analysis (PCA) of m go plant taxa relative abundances (%), indicating denser vegetation cover during the Holocene. For panels **a**, **b**, **c**, **d**, **e**, the top of each bar represents the median of samples within each 2000-year interval, while the lower and upper ends of the error bars indicate the first (Q1) and third (Q3) quartiles of those samples. Bars are plotted at the beginning of each interval. For panels **f**, **g**, the solid line indicates the median of six lake cores at 1000-year intervals, based on records interpolated to a common 1000-year grid. Timelines are indicated as Holocene, YD, Younger Dryas; B/A, Bølling-Allerød, Late Glacial; LGM, Late Glacial Maximum; MIS 3, Marine Isotope Stage.

The DNA weight percentages were converted using 10,000 iterations of a Monte Carlo simulation to DNA-projected organic carbon weight percentages (wt%) (OC_DNA-projected_), here defined as first order estimate of preserved sedimentary organic carbon inferred from sedimentary DNA. The conversion incorporated a range from maximum to minimum DNA C-values (Supplementary Fig. 4, 5), carbon content per cell, structural factors representative of each taxonomic group (listed in Supplementary Data 4), and extraction efficiency^36^. Additionally, a beta distribution was applied to microbial parameters to correct for cultivation biases in reference databases, which generally fail to reflect the reduced biomass and DNA-per-cell values characteristic of naturally occurring microorganisms^37, 38^(Supplementary Fig. 6). To ensure that dominant taxa are adequately represented, calculations were proportionally weighted using specific DNA and biomass-per-cell values of highly abundant plant and metazoa taxa (Supplementary Fig.7).

This resulted in a median OC_DNA-projected_ value of 1.2 wt% across all samples from all cores (Supplementary Data 3, range of sample-wise Monte Carlo medians: 0.0003–36.9 wt%), corresponding to roughly 1200 times the DNA wt% (Fig. 3b). Variations in median OC_DNA-projected_ across all lakes are shown in Fig. 3, while down-core changes for each individual lake are shown in Fig. 4b, revealing an overall increase toward the Holocene. The uncertainty of OC_DNA-projected_ (Fig. 4b) is shown for each sample as the interquartile range (IQR; Q1–Q3) derived from 10,000 Monte Carlo iterations (Supplementary Fig. 8). Given the large IQRs, OC_DNA-projected_ wt% values should be interpreted as approximations that provide a first-order rather than absolute estimate of carbon contributions.

**Figure 4.**
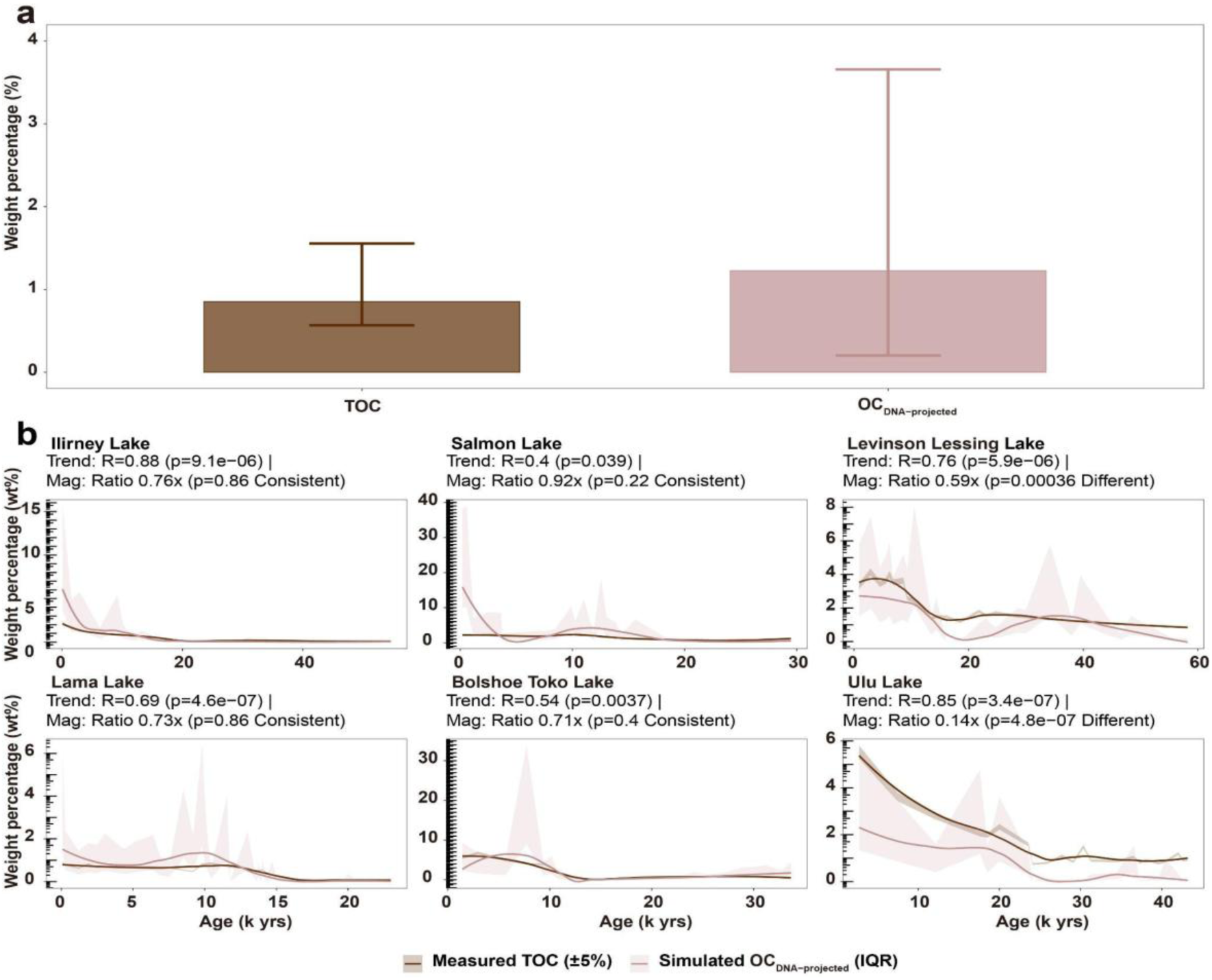
Comparison of OC_DNA-projected_ and biogeochemical organic carbon with source contributions per core and per age (k yrs). **(a)** Comparison of geochemical total organic carbon (TOC) with OC_DNA-projected_ weight percentage (wt%) across 161 samples from six lakes; the top of each bar represents the median of samples, while the lower and upper ends of the error bars indicate the first (Q1) and third (Q3) quartiles of those samples. **(b)** Temporal variations of OC_DNA-projected_ weight percentage (wt%, ±1 SD) are shown as shaded envelopes with LOESS-smoothed trends, alongside total organic carbon (TOC, ±5% uncertainty) over time.

### Comparison of OC_DNA-projected_ with TOC and evaluation of first-order controls on preserved sedimentary organic carbon

To evaluate our estimation of the preserved sedimentary OC pool, we compared OC_DNA-projected_ wt% with total organic carbon (TOC) wt%, measured mostly in the same samples (Fig. 4a, b). Overall, OC_DNA-projected_ wt% and TOC wt% are of similar magnitude across all samples (median OC_DNA-projected_ 1.2 wt%, median TOC: 0.85 wt%, Supplementary Data 3, Fig. 3b, c and 4a). More specifically, OC_DNA-projected_ and TOC show similar down-core variation across all lakes (R = 0.40–0.87, all p < 0.05, Fig. 4b). In terms of absolute values, agreement is strongest for Ilirney, Salmon, Lama, and Bolshoe Toko, whereas it is weaker for Levinson-Lessing and Ulu.

Given that TOC is widely regarded as a robust proxy for organic matter content in lake sediments^39^, our inference of similar magnitudes and mostly strong correlations with TOC underlines the reliability of our estimations of OC_DNA-projected_. Considering that the DNA content per cell (DNA-C values) and the taxon-specific biomass per cell differ by several orders of magnitude among the taxa^40, 41^, and given the high uncertainties in the measurement of DNA content, the quantitative and trend agreement between the two proxies is remarkable. By employing a beta distribution focused on the lower spectrum of genome size and cellular biomass, we aim to provide more realistic estimates than those derived from ideal laboratory settings. Nevertheless, we acknowledge that this approach might not capture the full range of variability driven by the life stages or environmental stressors of specific microorganisms. However, as our OC_DNA-Projected_ weight percentages show a strong fit with the bulk TOC record, we hypothesize that the impact of this missing variability on the overall carbon budget remains minimal.

Our finding that OC_DNA-projected_ wt% and TOC wt% show similar down-core trends and, in most lakes, similar magnitudes indicates that sedimentary DNA and bulk organic carbon were shaped by broadly similar depositional and stabilization controls. We therefore interpret the taxonomic signal underlying OC_DNA-projected_ as a first-order, semi-quantitative proxy for the magnitude and taxonomic composition of the preserved sedimentary OC pool, rather than as evidence that DNA and bulk organic matter have identical decay kinetics. DNA is intrinsically more labile than structural biopolymers such as lignin or algaenan^42^, but its persistence can be extended by physical shielding. In lake sediments, green-algal and vascular-plant DNA can decline at rates similar to corresponding lipids and lignin when protected within recalcitrant cell walls^6^. Extracellular DNA can also be stabilized after burial by adsorption to mineral surfaces, which reduces nuclease degradation^43^, and in marine sediments it is commonly associated with mineral matrices and organic–inorganic aggregates rather than occurring only inside living cells^28^. Bulk organic matter is governed by partly analogous controls, because oxygen exposure time regulates burial efficiency in lake sediments^5^, and mineral-bound organic carbon can persist for millennia relative to unbound organic carbon^44^.

Taken together, these studies suggest that, after strong early-diagenetic filtering, stabilized fractions of DNA and bulk organic matter can persist on similar millennial timescales, so differential degradation is unlikely to be the dominant first-order control on the down-core signal. This interpretation is probably weakest in the youngest sediments, where fresh extracellular DNA, intact cells, or viable resting stages may still contribute substantially^45^. It is also consistent with a two-stage model of DNA persistence, with rapid early loss followed by much slower residual decay^46, 47^. This may explain why the youngest samples in most cores show relatively high OC_DNA-projected_ values and little late-Holocene decline. By contrast, the TOC excess in Levinson Lessing and Ulu is more plausibly explained by additional inputs of age (Fig. 4b), DNA-poor allochthonous carbon from large catchments than by enhanced DNA preservation alone, which is also consistent with the abundant black particles in the pollen slides^48^. Future versions of genC could therefore distinguish a young, labile DNA-rich pool from an older, mineral-associated stabilized pool, constrained by sediment age, DNA damage, and mineralogical and depositional variables that influence both DNA preservation and long-term OC burial. In addition, while we minimized potential methodological biases through stringent read quality control and end-to-end alignment, residual GC-related recovery bias between groups may still influence read abundance estimates and should therefore be considered a limitation when making quantitative comparisons across groups.

Despite the potential for site-, time- and taxon-specific offsets, our observations suggest that sedaDNA-derived OC_DNA-projected_ wt% may represent a novel semi-quantitative proxy for assessing first-order variations in the magnitude and composition of organic matter preserved in lake sediments over millennial to glacial–interglacial timescales. This is consistent with previous studies showing that DNA itself can serve as an indicator of organic carbon content, exhibiting a positive linear relationship with organic carbon in soils and lake sediments^49, 50^. Furthermore, DNA-based reconstruction of carbon contributions in a marine context has provided a more complete picture of terrigenous organic matter burial than DNA read composition along^20^. Unlike traditional methods that rely on plant-group-specific biomarkers such as lignin^16^, leaf waxes^51^, or taraxerol^52^, DNA captures molecular traces from a broad range of biomass sources, including major contributors such as roots and leaves, directly within the sediment, thereby complementing the existing biogeochemical proxy ensemble.

### OC_DNA-projected_ burial rates and environmental and preservation control

We used sample-specific sedimentation-rate estimates derived from age–depth models to quantify total OC_DNA-projected_ burial rates for the individual sediment records. These ranged from 0.000001 to 0.03 g cm^−2^ year^−1^, with a median of 0.00031 g cm^−2^ year^−1^ (Supplementary Data 3, Fig. 3d). Our results show that OC_DNA-projected_ burial rates are of a similar order of magnitude to long-term OC burial rates reported from extra-tropical regions with broadly comparable environmental settings. For example, Ferland et al.^53^ reported a Holocene OC burial rate of approximately 0.00038 g cm^−2^ year^−1^ for lakes in northern Québec, Canada. Similarly, Kortelainen et al.^54^ estimated an average Holocene burial rate of 0.00018 g cm^−2^ year^−1^ based on data from 122 lakes across Finland. Overall, we found intermediate median burial rates for MIS 3 (60–26 ka, median: 0.00031 g cm^−2^ year^−1^), low median burial rates during MIS 2 (26–12 ka, median: 0.0001 g cm^−2^ year^−1^), and high values during the Holocene (<12 ka; median: 0.00079 g cm^−2^ year^−1^) (Figs. 2c and 3d). Likewise, Li et al.^55^ reported an increase in lake carbon burial rates from 0.00023 g cm^−2^ year^−1^ during the Last Glacial Maximum to 0.00064 g cm^−2^ year^−1^ during the mid-Holocene based on lakes in northwest China.

A generalized linear mixed model (GLMM) analysis evaluating potential drivers of OC_DNA-projected_ burial rates (Supplementary Fig. 9, Table 1) revealed that higher OC_DNA-projected_ burial rates were significantly associated (p < 0.1) with higher sedimentation rates, higher temperatures, and denser vegetation cover, represented by higher PC1 scores of plant DNA composition (Fig. 3 f, g, Supplementary Fig. 10, 11, 12). These findings agree with previous studies that likewise identified warmer temperatures and more developed vegetation as favorable conditions for higher organic matter content and burial rates in lake sediments ^8^. This relationship is most likely explained by greater nutrient availability, particularly nitrogen and phosphorus, fueling primary productivity within both lake systems and their catchments ^56^.

**Table 1.** Summary of coefficients estimated by Gaussian generalized linear mixed models (GLMMs) to evaluate the effects of predictor variables on the Burial rate OC_DNA-projected_ weight percentage.

| Predictor variables | Total burial rate OC<sub>DNA</sub>-projected |  | Percentage of aquatic OC<sub>DNA</sub>-projected weight percentage |  |
| --- | --- | --- | --- | --- |
|  | Estimate | *P*-value | Estimate | *P*-value |
| Age | -0.11462 | 0.45037 | **-5.1238** | **6.17e-05 \*\*\*** |
| Average read length (bp) | 0.07946 | 0.37635 | 1.3011 | 0.1592 |
| Sedimentation rate | **0.37242** | **0.00340 \*\*** | **-2.1864** | **0.0497 \*** |
| Dense vegetation (PC1) | **0.44714** | **0.00126 \*\*** | **2.6409** | **0.0426 \*** |
| Annual precipitation (P<sub>ann</sub>) | 0.02085 | 0.85668 | -1.2576 | 0.2050 |
| July temperature (T<sub>Jul</sub>) | **0.20814** | **0.06523 .** | -1.3386 | 0.2268 |

*P*-value: two-sided t-tests using Satterthwaite's Predictor variables with *P*-value < 0.1 is in bold Significance codes: \*\*\* 0.001; \*\* 0.01; \*0.05; . 0.1.

Sediment age showed a weak negative relationship with OC_DNA-projected_ burial rates, whereas average read length showed a weak positive relationship (Table 1). These effects have the expected sign and may reflect degradation^57^, but they are weak and do not appear to represent first-order controls on OC_DNA-projected_ burial rates compared with the climatic and depositional drivers of organic carbon burial.

In contrast to the temporal within-lake relationships, OC_DNA-projected_ burial rates were generally lower in boreal lakes than in Arctic tundra lakes (Fig. 2b), suggesting that spatial and temporal gradients likely reflect partly different ecological and environmental drivers. However, we currently cannot provide a direct explanation for this difference as DNA preservation signals, reflected by read fragment length (Supplementary Fig. 13), as well as depositional setting (Supplementary Table 1), do not differ substantially between these tundra lakes and the others.

### Temporal variation in aquatic and terrestrial sources of organic carbon in lake sediments

Lake sediments contain organic matter from both aquatic and terrestrial sources^21^. To better resolve the sources of the preserved OC pool, we assigned sequencing reads to aquatic or terrestrial habitats using a continuous affinity framework. Habitat assignment was carried out at the species level for microorganisms (Bacteria and Archaea) and at the family level for Eukaryota. For Bacteria and Archaea, we developed and applied MICROSPyDER, a custom web crawler that systematically retrieves structured habitat metadata from the MicrobeAtlas database (https://microbeatlas.org/), whereas Eukaryota were classified using literature-based habitat associations (Fig. 1, Supplementary Data 5). Across all lakes, 58.5% of reads met these classification criteria. Within this fraction, a median of 4.5% of reads was assigned to aquatic organisms, 71.0% to terrestrial organisms, and 16.2% remained unclassified with respect to habitat.

Our source separation indicates that terrestrial sources dominate the preserved sedimentary OC pool, with a smaller contribution from aquatic sources (Fig. 5b, Supplementary Data 3, terrestrial: 0.89 wt% (Range: 0.0021-37.16 wt%), percentage of terrestrial OC_DNA-projected_ weight percentage: 95.2%; aquatic: 0.07 wt% (Range 0.00026-9.81 wt%), percentage of aquatic OC_DNA-projected_ weight percentage: 4.8 %; medians across all samples). Given the relatively large size of these lakes, this substantial terrestrial contribution may seem surprising, yet previous studies have likewise emphasized the importance of terrestrial organic carbon inputs for lake sediment carbon stocks^22^.

**Figure 5.**
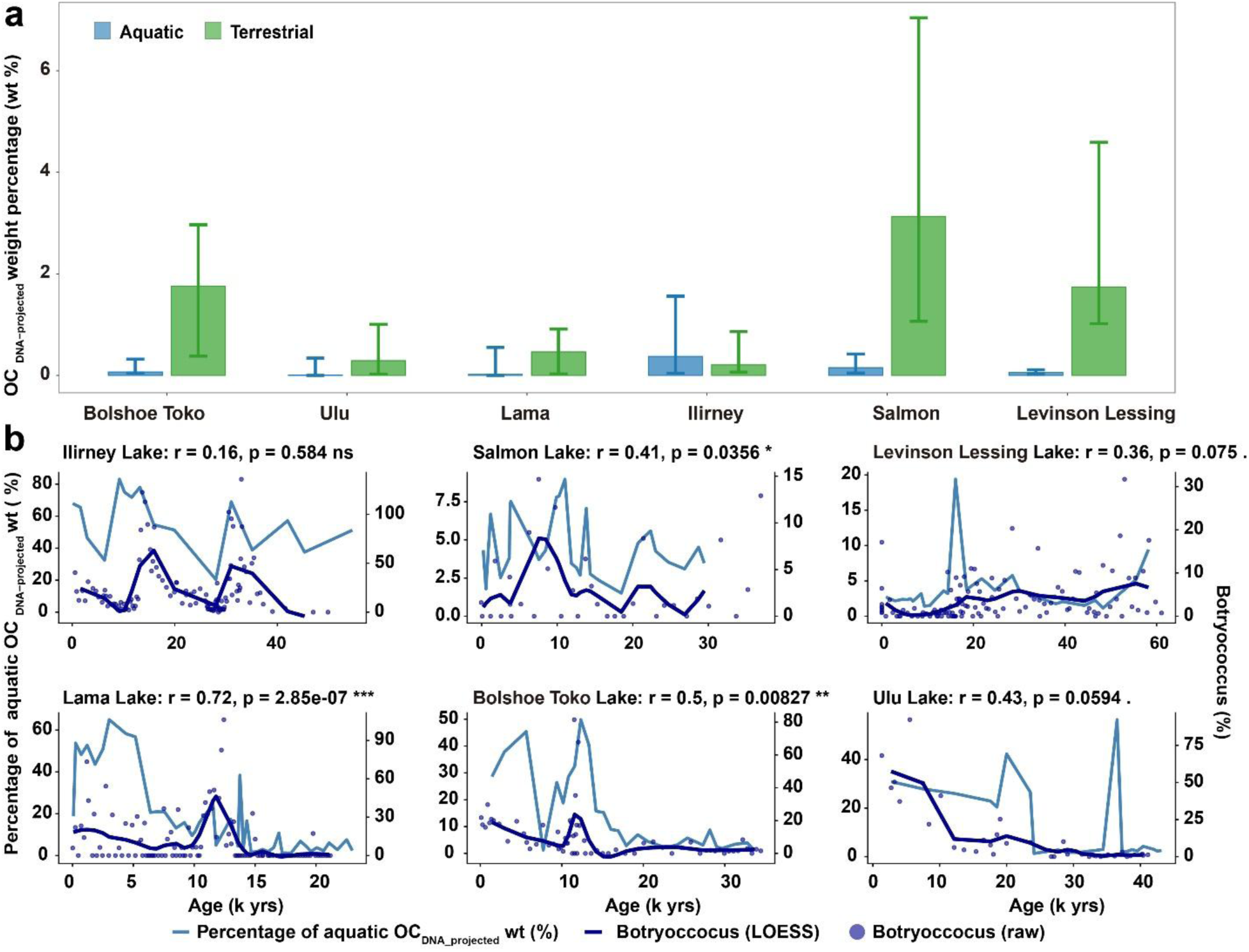
Aquatic and terrestrial contributions to sedimentary OC_DNA-projected_ and their relationship with *Botryococcus* abundance across six lakes. **(A)** Comparison of the relative contributions of aquatic and terrestrial sources to OC_DNA-projected_ weight percentages (wt%) across six lakes; Aquatic OC wt is shown in blue and terrestrial OC wt in green; the top of each bar represents the median of samples, while the lower and upper ends of the error bars indicate the first (Q1) and third (Q3) quartiles of those samples. **(b)** Each lake is shown with percentage of aquatic OC _DNA-projected_ weight percentage (%, left axis) and Botryococcus percentage (%, right axis). Raw Botryococcus values are displayed as dark blue points, and LOESS-smoothed trends (span = 0.3) as dark blue lines. The right y-axis shows Botryococcus percentages, which were linearly rescaled to match the range of aquatic biomass percentages for visualization purposes. Correlation analyses were performed using the original (unscaled) data. Spearman’s rank correlation coefficients (r) and associated significance levels are provided within each panel (. P < 0.1; * P < 0.05; ** P < 0.01; *** P < 0.005).

We evaluated the temporal pattern of reconstructed aquatic contribution (expressed as percentage of total OC_DNA-projected_ wt%) using *Botryococcus* abundance, expressed as a percentage of the terrestrial pollen sum, as a fully independent proxy. *Botryococcus* is a colony-forming freshwater green alga that commonly occurs in high abundances in lacustrine palynological records and is widely used as a semiquantitative indicator of open-water conditions and aquatic productivity^58, 59, 60, 61^. *Botryococcus* abundance was positively correlated with reconstructed aquatic percentages across all six cores (Fig. 5b, Spearman ρ = 0.16–0.72); correlations were significant in most lakes (p < 0.1), but not in Lake Ilirney (ρ = 0.16, p = 0.58). These results generally support the validity of our reconstruction of aquatic contributions, while also highlighting some lake-specific variability in proxy performance. Although OC_DNA-projected_ wt% are low during glacial periods, the relationship with *Botryococcus* remains consistent. However, large percentage shifts may have limited ecological significance in absolute terms when total carbon is very low.

We refrain from using bulk C/N values as a simple terrestrial–aquatic source proxy for evaluating aquatic contributions to OC_DNA-projected_ wt%, because in permafrost catchments the major endmembers are not clearly separated. Degraded permafrost-soil organic matter^62^, aquatic and terrestrial microbial biomass, and aquatic algae^10, 63^ can all show relatively low C/N values within the range observed in our sediments (mostly 5–15)^57, 64, 65, 66, 67, 68^.

A GLMM indicates that the percentage of aquatic OC_DNA-projected_ wt% increased toward the present and with denser vegetation, but decreased with higher sedimentation rate (Table 1). Both changes in original input and preferential preservation may have contributed to this pattern. The percentage of aquatic OC_DNA-projected_ wt% increase from the glacial to the Holocene (Fig. 3e), although some lake records deviate from this median trend (Fig. 5b). Warming during the early Holocene, together with longer ice-free seasons, enhanced nutrient input from more developed terrestrial ecosystems, and reduced turbidity following the disappearance of direct glacier input, may have increased lake-internal productivity relatively more than terrestrial productivity^69^. This interpretation is further supported by spatial patterns showing that Holocene values in Salmon Lake and Lake Levinson-Lessing, located in the Arctic tundra zone, display the lowest proportions of aquatic OC_DNA-projected_ wt% compared with lakes from the boreal zone (Fig. 5a). In addition, Holocene development of permafrost wetlands^70^ in lowland areas surrounding the lakes presumably increased the influx of allochthonous aquatic organic matter^71^.

At the same time, preferential degradation of aquatic organic matter may also have influenced the reconstructed pattern. Aquatic organic matter is typically more nutrient-rich and less structurally complex than terrestrial organic matter^21^, making it more labile and more prone to early diagenetic degradation, which may have contributed to preferential preservation of terrestrial material. This may have led to an underestimation of the aquatic contribution at the time of sedimentation particularly in older sediments^63^. It may also help explain why age emerged as a stronger predictor of aquatic contributions than, for example, temperature or vegetation, which were also relatively high during MIS 3 but did not result in similarly high reconstructed aquatic contributions.

### Contribution of different taxon groups to lake organic carbon over time

Building on the distinction between aquatic and terrestrial sources of preserved sedimentary carbon, taxon-level estimations provide a more detailed picture of specific carbon sources and their temporal variability.

To assess the impact of parameter-specific uncertainties in translating DNA into OC_DNA-projected_ estimates, we performed a Sobol sensitivity analysis and a perturbation analysis in which key parameters (cellular biomass, genome size, DNA extraction efficiency, and structural assumptions) were varied from 50–90% to 110–200% of their baseline values. The sensitivity analyses show that OC_DNA-projected_ estimates are primarily driven by genome size and cellular biomass, with two-fold perturbations causing approximately ±100% changes in median values, whereas uncertainties related to DNA extraction efficiency and structural factors have smaller effects (<30%). Although the absolute OC_DNA-projected_ estimates changed, the relative ranking of major taxonomic groups remained largely stable, with dominant contributors being consistently identified across all scenarios, supporting the robustness of the GenC approach to uncertainty in single-parameter choices (Supplementary Fig. 14, 15).

Our results indicate that OC_DNA-projected_ (wt%) vary much more widely among taxa than the corresponding DNA weight percentages (Fig. 6a). Aquatic algae/protists (∼2,522-fold) show the highest amplification relative to their DNA wt%, reflecting the high variability in DNA weight per cell^72^. Woody plants (∼1,040-fold) and non-woody plants (∼1,020-fold) also show high amplification. Viridiplantae and fungi exhibit approximately 450-fold and 270-fold amplification, respectively. Archaea, bacteria, and metazoa show about 80-fold, 55-fold, and 30-fold amplification, whereas viruses display only minimal amplification (∼2.7-fold).

**Figure 6.**
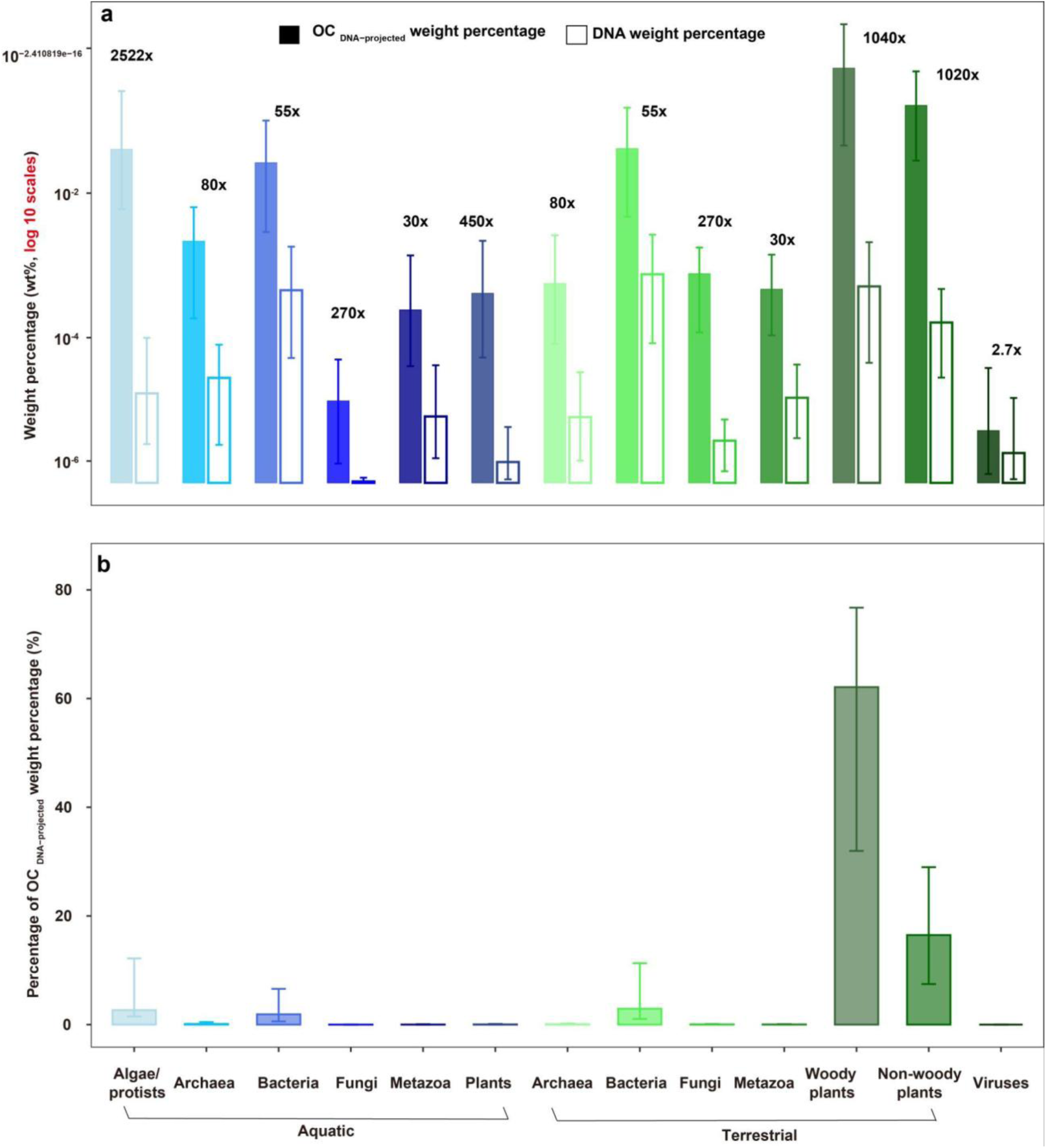
Weight percentage Content and relative OC_DNA-projected_ weight percentage abundance per organism groups. **(a)** Weight percentage (wt%) of OC_DNA-projected_ (solid bars) and DNA (hollow bars) across aquatic and terrestrial per organism group on a log10 scale; **(b)** Percentage contribution of OC_DNA-projected_ weight percentage (wt%) per organism group. The top of each bar represents the median of samples, while the lower and upper ends of the error bars indicate the first (Q1) and third (Q3) quartiles of those samples. Aquatic organism groups displayed on the left and terrestrial organism groups on the right. For panels **a**, the projection ratio was calculated as the ratio between the median OC_DNA-projected_ weight percentage and the median DNA weight percentage across 161 samples from six lakes within each organism group.

**Figure 7.**
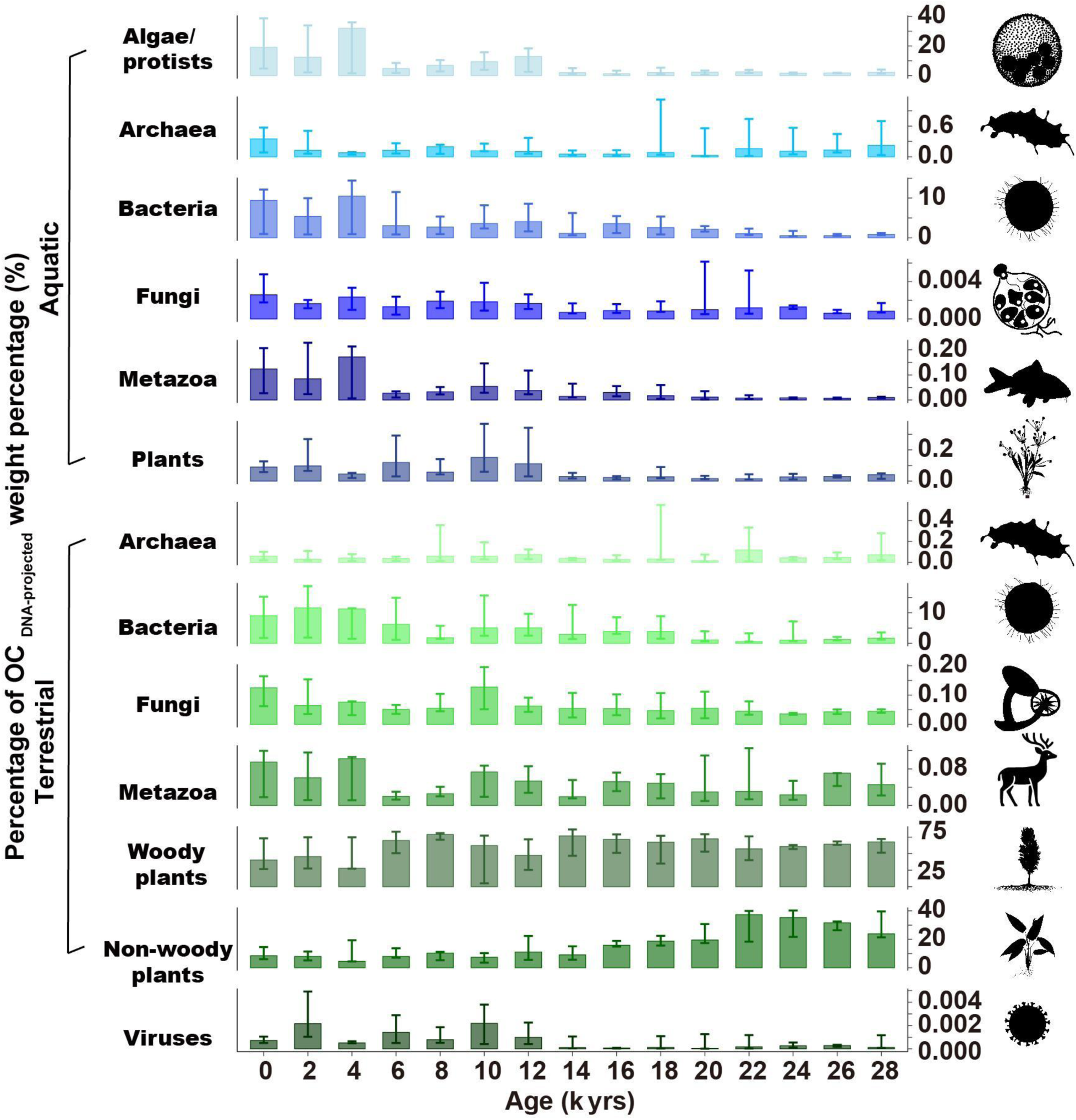
Percentage of OC_DNA-projected_ weight percentage (%) for organism groups in six lake cores over the past 30,000 years. The taxonomic groups are separated by the habitat they originate from. The top of each bar represents the median of samples within each 2000-year interval, while the lower and upper ends of the error bars indicate the first (Q1) and third (Q3) quartiles of those samples. Aquatic organism groups are shown in the upper panels, and terrestrial organism groups in the lower panels. Silhouette images are provided under the CC0 1.0 Universal license. Arrows indicate the direction of temporal trends inferred from the fitted Hierarchical Generalized Additive Models (HGAMs)

These patterns highlight the need to account for taxon-group-specific parameters when aiming for quantitative reconstruction of taxon-specific contributions to the preserved OC pool and, in turn, for assessing carbon burial. Without such corrections, key carbon contributors—especially aquatic algae/protists, woody plants, and non-woody plants—may be severely underestimated as sedimentary carbon sources.

Our data reaffirm the central role of vascular plants as major terrestrial contributors to the organic carbon (OC) pools preserved in lake sediments, as has also been shown for marine sediments^20^. Based on the median values across all samples, terrestrial plants accounted for 80% of the total OC_DNA-projected_ OC. Woody plants alone contributed 60%, indicating that they contribute disproportionately more to preserved sedimentary carbon burial than non-woody plants (Fig. 6b). The abundance of recalcitrant cell-wall components, such as lignin, may physically protect DNA and other labile biomolecules from microbial degradation, thereby enhancing long-term preservation in lake sediments^6^. Although relative contributions from non-woody plants peaked during the cold MIS 2 (Fig. 7, Supplementary Fig. 16), the contribution of woody taxa remained high throughout the glacial period (Fig. 7, Supplementary Fig. 17). This may seem counterintuitive given that tundra-steppe dominated Beringia during this period^73^ (see PCA analyses of plant composition, Supplementary Fig. 10). However, dwarf shrubs such as *Salix* and *Dryas* were common elements at our sites throughout the glacial period^74^ (Supplementary Fig. 17), consistent with regional reconstructions^75^. In addition, we assume that woody taxa are disproportionately represented because of their extensive root systems^76^ and preferential establishment along watercourses, particularly *Salix*^20^ (Fig. 7, Supplementary Fig. 16). Furthermore, the absolute burial of woody biomass was significantly higher during the Holocene than during the glacial period (Supplementary Fig. 18), likely reflecting generally higher terrestrial plant productivity in the more complex ecosystems that developed under warmer conditions^33, 77^.

Our results indicate that aquatic and terrestrial metazoans contributed only marginally to OC_DNA-projected_, with a median of 0.05% of total wt% (Fig. 5b, Supplementary Data 3), and remained minor even during the megafauna-rich Pleistocene^78^. This is roughly an order of magnitude lower than the contribution of animals to modern global biomass^79^. A likely explanation is that nutrient-rich metazoan biomass is strongly recycled in food webs before burial, as supported by fish sedimentary eDNA studies^80^, and that its protein-rich organic matter is preferentially degraded in sediments^81^. In addition, taxonomic database bias, particularly for arthropods, may have led to further underrepresentation of metazoan inputs.

Overall, aquatic algae/protists form a substantial part of the lake-sediment carbon budget. However, only with the onset of the Holocene did median primary production by eukaryotic algae rise above 10% (Fig. 6, Supplementary Fig. 16). Strong physical mixing and increased turbidity caused by meltwater, and the related reduction in light availability for photosynthesis, may have constrained the growth of eukaryotic algae^82^. In contrast, the contribution of aquatic bacteria increased already during the deglacial (Fig. 6).

Aquatic and terrestrial bacteria emerge in our data as the strongest contributors to the preserved OC pool in lake sediments, aside from terrestrial plants. Median terrestrial bacterial OC_DNA-projected_ wt% accounts for 3% of the total OC wt%, while aquatic bacterial OC_DNA-projected_ wt% accounts for 2% (Fig. 6b, Supplementary Data 3), consistent with previous studies showing that organic matter derived from bacterial biomass can constitute an important fraction of sedimentary organic matter (0.01–43.6%^83^). The organic matter produced by bacteria is rich in complex, recalcitrant biomolecules that can persist in sediments over millennia^84^. This interpretation is further supported by our observation of a C-to-T exchange pattern in bacteria, with 50–70% of bacterial reads mapping to damaged contigs, which is in a similar range as in other taxonomic groups and suggests a predominantly ancient origin (Supplementary Fig. 1; see also Herzschuh et al.^20^). Supporting this view, previous studies found that most bacterially derived organic matter in permafrost soils^85^ and lake sediments^86^ is necromass. Our results suggest that bacterial organic matter from both aquatic and terrestrial sources can be stabilized and retained over glacial–interglacial timescales, raising the possibility that bacteria play a more substantial role in long-term carbon sequestration than previously appreciated.

Among microbial groups, Archaea, although minor contributors overall (median: 0.2% of total OC_DNA-projected_ wt%, Supplementary Data 3), show a slightly higher relative OC_DNA-projected_ signal during the cold glacial period than during the Holocene (Fig.7, Supplementary Fig. 16), consistent with their known preference for extreme environments such as permafrost soils^87^.

Fungi OC_DNA-projected_ contributions are low relative to bacterial contributions in our dataset (Fig. 6b, 7). However, it is inline with the idea that fungi may contribute more strongly to short-term carbon turnover than to long-term preserved OC pools^88^ which may explain the high values in the youngest time-slice (Fig. 7, Supplementary Fig. 16). Our results show that relative and absolute terrestrial fungal contributions increased at the transition to the Holocene when the modern soil regime became established at the sites along with forest expansion^23^ (Supplementary Fig. 18). Because relative DNA signals can be influenced by recovery and reference biases, we interpret this pattern cautiously.

Likewise, viruses contribute least to the OC_DNA-projected_ due to their minimal carbon content (Fig. 6)^89^. A virus-Host study of this data set revealed that most DNA viruses have bacterial and algal hosts and show a positive co-variation with them^90^. This is in line with the finding that viruses along with the major host group OC_DNA-projected_ increase with the Holocene (Fig.7, Supplementary Fig. 16). We considered potential influences of extraction bias, GC content, and fragment length, but interpretation of these patterns should still account for inherent uncertainties associated with read-based and reference-dependent approaches.

## Conclusions

We developed the genC pipeline to translate sedimentary ancient DNA into OC_DNA-projected_, a first-order, semi-quantitative estimate of taxon-resolved contributions to preserved organic carbon in lake sediments. Applied to six lake records from Siberia and Alaska spanning the last 30,000 years, OC_DNA-projected_ broadly reproduced independent patterns in total organic carbon and aquatic contribution, supporting its use as a complementary proxy for long-term changes in the preserved sedimentary OC pool.

Across the records, preserved OC was dominated by terrestrial inputs, especially woody plants, whereas aquatic algal contributions generally increased during the warmer Holocene; bacterial-derived organic matter formed a persistent additional component, while metazoan contributions remained minor. Together, these results suggest that millennial-scale variation in preserved OC was driven mainly by environmental change, particularly warming and vegetation development, rather than by preservation alone. Although absolute estimates remain uncertain because of preservation effects and taxon-specific trait variability, genC provides a new taxonomically resolved framework for linking palaeoecological change with long-term carbon burial in lakes.

## Methods

### Sediment Cores

Six lake sediment cores were analyzed for this study, five from Siberia and one from Alaska. The Siberian cores are Ilirney (67.35°N, 168.32°E), for which detailed descriptions and the age-depth model can be found in Vyse et al.^68^; Ulu (63.34°N, 141.04°E), with its age-depth model published in Jia et al.^57^; Bolshoe Toko (56.25°N, 130.50°E) described in Courtin et al.^65^; Levinson Lessing (74.47°N, 98.67°E) documented in Scheidt et al.^66^; and Lama Lake (69.53°N, 90.20°E) with details available in von Hippel et al.^67^ and Andreev et al.^64^. The Alaskan core, Salmon (64.91°N, 164.99°W), is also included in this study, with its dating and age-depth model to be presented in PANGAEA^91^.

### Extraction of sedaDNA and shotgun metagenomics

This study uses a shotgun dataset published by Von Hippel et al.^23^ for Lake Lama and Liu et al.^33^ for the other 5 cores. The sedaDNA extraction and shotgun sequencing of the six sediment cores are described in detail in Liu et al.^33^. Briefly, all sediment cores were subsampled in a clean climate-controlled chamber at GFZ Potsdam, Germany. DNA extraction and single-stranded library preparations were carried out in the palaeogenetic laboratories at Alfred Wegener Institute (AWI) Helmholtz Centre for Polar and Marine Research, Potsdam, Germany. DNA was extracted from 161 sediment samples using the DNeasy PowerMax Soil Kit (Qiagen), followed by purification and concentration with the GeneJET PCR Purification Kit (Thermo Fisher Scientific). DNA extraction blanks (EB) were run with each extraction batch generally consisting of nine sediment samples (from one sediment core) and one extraction blank (control of extraction chemicals and extraction procedure). Finally, DNA from samples and blanks was quantified with the Qubit Fluorometer (Thermo Fisher Scientific) and adjusted to a concentration of 3 ng µL^−1^ per sample The concentration of the extraction blanks ranged between 0 to 0.2 ng µL^−1^. For some lake cores extraction blanks were pooled prior to library preparation and sequencing (see Supplementary Data 2). 15 to 30 ng of the DNA extracts of the samples and 3 µL of the extraction blanks were used as input material to the single-stranded DNA library preparation following the protocols of Gansauge and Meyer^92^ and Gansauge et al.^93^ with modifications described in^22^. A library blank (LB, only library chemicals) was run for each library batch. Double indexed DNA libraries from samples were pooled in equimolar amounts, while 3 µL of the extraction and library blanks were added. Samples and blanks (EB & LB) from each sediment core were sequenced in-house on the Illumina NextSeq 2000 device at AWI, Bremerhaven, Germany, by using P3 Reagents and a customized sequencing primer (CL72) for 2×100bp paired-end (200 cycles) sequencing, except for Lake Ilirney and some Lama Lake samples which were sequenced at 2×100bp paired-end mode at the company Genesupport Fasteris (Switzerland) using the Illumina NextSeq2000 and NovaSeq 6000 device, respectively. Raw sequence data of the six cores are published in the European Nucleotide Archive for accession number see Data and code availability.

### Bioinformatic analyses and sedaDNA data quality control

The taxonomic classification of raw sequencing data from samples and blanks followed the HOLI pipeline^94^ with modifications described in Liu et al.^22^ and expansion of the reference database by including virus genomes^33^. Reads were quality-filtered and deduplicated, and filtered to a minimum read length of 30 bp. Reads were aligned in end-to-end mode with Bowtie2^95^ to the reference database and subjected to LCA-based taxonomic assignment using the ngsLCA algorithm^96^ (implemented via metaDMG^97^), applying a minimum identity threshold of 0.95. Up to 1000 valid and unique alignments per read were retained (-k 1000). Across the six lakes, a total of 64 blanks were processed (Ilirney lake: 5, Salmon lake: 14, Levinson Lessing lake: 10, Lama lake: 19, Bolshoe Toko lake: 5, Ulu lake: 11) (Supplementary Data 2). Therefore, taxonomically classified reads from samples and blanks were aggregated into read counts per taxon per sample and sample metadata (depth and age of the samples and the blanks) were merged and combined to a final data file, which was further analysed in R version 4.4.2^98^.

Post-mortem damage patterns were investigated with PyDamage v0.72 (Borry et al.^99^; as described in von Hippel et al.^23^ and Çabuk et al.^100^). In short, quality-filtered reads from samples and blanks were error-corrected with BBtools v38.87^101^ and de novo assembled into contigs using MEGAHIT v1.2.9^102^. Due to the low number of read counts in the extraction and library blanks the assembly step failed in almost all blanks supporting the cleanliness of the laboratory procedure. Assemblies resulting from the samples were then further analyzed in pyDamage and the final alignments were used as input to estimate the post-mortem damage of the contigs, which were subsequently taxonomically classified using Kraken2 against the database (downloaded in October 2022) with a confidence threshold of 0.0. PyDamage output was investigated for all lake sites and seven taxon groups: bacteria, archaea, viruses, fungi, plants (Viridiplantae), metazoa (excluding reads/contigs from Hominidae), and algae that were filtered with a taxa based on the HOLI taxonomic classifications using the best taxonomic level (column “taxon” in the HOLI output). Contigs for each lake core classified to the defined seven taxon groups were filtered for a prediction accuracy ≥ 0.5 and a contig length ≥ 500 bp to identify ancient contigs. C-to-T substitution frequency of the first 10 read positions of the 5’end was plotted for all lake sites and the seven taxon groups aggregated over all sample ages. The proportion of damaged reads was calculated for each taxa group and site. The total number of reads for each taxa group and site was calculated and the number of reads after using the filter criteria prediction accuracy ≥ 0.5 and a contig length ≥ 500 bp. The proportion of filtered reads from total is the % of damaged reads. Sample-wise assessment was possible for Lake Lama only that was sequenced with higher effort. The relationship between C-to-T substitution frequency of the first read position across sample age per each taxa group using Pearson correlation.

### Dataset preparation

All statistical analyses and data visualization were performed in R Studio (R version 4.4.2^98^). The initial data processing involved importing and structuring ancient DNA (aDNA) datasets from multiple files using the R packages *data.table*^103^, *stringr*^104^, and *tidyverse*^105^.

### Data mining and automated habitat assignment

For the analysis, only DNA sequence reads taxonomically classified to at least to Superkingdom (Bacteria, Archaea, Viruses, Eukaryota) level were considered. Eukaryotic reads were further classified at the Kingdom level and grouped into Fungi, Metazoa, Viridiplantae, and aquatic algae/protists. This category includes a broad range of predominantly aquatic eukaryotic phyla detected in our dataset, encompassing both photosynthetic groups (Bacillariophyta, Rhodophyta, Rhodophyta), and heterotrophic or non-photosynthetic protistan groups (Discosea, Oomycota, Evosea, Euglenozoa, Tubulinea, Cercozoa, Apicomplexa, Heterolobosea, Ciliophora, Fornicata, Parabasalia, Preaxostyla, Nebulidia, Endomyxa, Picozoa, Foraminifera, and Perkinsozoa).

To address the challenges of high-resolution habitat classification for all Bacteria and Archaea classified on species level, we developed MICROSPyDER (MICRObial Source crawling & Integration Python Data Engine for Relative-affinities). This integrated pipeline replaces traditional discrete classification with a continuous affinity spectrum, utilizing a custom web crawler to systematically retrieve structured metadata from the MicrobeAtlas database (https://microbeatlas.org/).

Implemented in Python (python v3.12), the MICROSPyDER framework employs the *webdriver* function (Selenium) for browser automation and BeautifulSoup4 for parsing dynamic HTML content. The engine programmatically queried each bacterial and archaeal strain on species level, utilizing regular expressions (*re*) for pattern matching and pandas for data integration. For each taxon, the crawler extracted the relative frequency (*V*) across diverse environmental categories, including primary ecological tiers (e.g., Aquatic, Terrestrial) and their nested sub-habitats (e.g., Marine, Freshwater, Soil). To ensure data integrity and prevent server overload, the pipeline included explicit wait conditions and temporal delays (1–10 seconds). Taxa yielding no direct matches were flagged for manual verification.

The core of the MICROSPyDER pipeline is the calculation of a Habitat Affinity Score (HAS), which enables the placement of species along a continuous habitat spectrum. This score ( *H*_*i*_ ) contrasts the frequency of a taxon *i* in terrestrial versus aquatic environments:

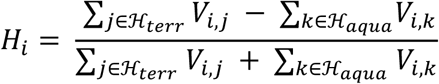

In this equation, *V*_*i*,*j*_ and *V*_*i*,*k*_ represent the observed frequency of taxon *i* within a specific habitat. The symbol **∈** denotes the membership of a specific habitat category within a predefined set; thus, *j* ∈ ℋ_*terr*_ refers to all habitat categories *j* that are elements of the terrestrial set (ℋ_*terr*_), while *k* ∈ ℋ_*aqua*_refers to all categories *k* belonging to the aquatic set (ℋ_*aqua*_).

The score is normalized by subtracting the sum of aquatic frequencies from the terrestrial frequencies and divided by the total sum of detections. This allows for a robust habitat classification where the resulting HAS values per species range from aquatic specialists (−1), through generalists (0), to terrestrial specialists (1) per core (Supplementary Fig. 19–24). Bacterial and archaeal species were assigned to one of the two habitats in this manuscript when they reached at least a HAS score of 0.5 for terrestrial or -0.5, for aquatic respectively. Species with HAS values between −0.5 and 0.5 had their reads proportionally partitioned between aquatic and terrestrial fractions using linear interpolation across the threshold interval. Specifically, the fraction assigned to each habitat was determined by the relative position of the HAS value within the interval, such that taxa closer to the aquatic threshold contributed a larger fraction of reads to the aquatic pool, whereas taxa closer to the terrestrial threshold contributed a larger fraction to the terrestrial pool.

For Metazoa, Viridiplantae and Fungi we used automated literature research at family level, by applying the *regex* package in RStudio (R version 4.4.2^98^) to determine the origin habitat and in addition manually checked the results at family level because habitat preferences are generally conserved within families in these groups^106^. Although metagenomic classification may be affected by database bias, the detected taxa (e.g., *Betula*, Ericaceae, Salicaceae, Rosaceae) are consistent with expected regional vegetation, supporting the robustness of our assignments^20^. Terrestrial Viridiplantae were additionally subdivided into woody and non-woody groups. Due to the ecological complexity, low biomass and broad host ranges of viruses, habitat-based classification for viral taxa was not attempted^90^.

Reads not assigned to any taxon were first distributed to known groups in proportion to the abundances of already classified reads within each group. They were then combined with unclassified habitat reads and reads only classified to superkingdom or kingdom level and subsequently distributed into aquatic and terrestrial fractions according to the relative abundances of already classified aquatic and terrestrial reads within each group and sample.

### Organic carbon contribution calculation - the genC pipeline

The genC pipeline integrates taxonomically assigned DNA data with geochemical bulk parameters—such as sedimentation rates and dry bulk density. By factoring in taxon-specific biomass and nuclear DNA content (pg) (DNA C-value) as well as structural factors (e.g., carbon-rich tissues and compounds without DNA, such as xylem or extracellular polymeric substances) and extraction efficiency, it enables a quantitative assessment of each taxon’s individual contribution to the total sedimentary carbon pool.

OC_DNA-projected_ was calculated for the major taxonomic groups separated into aquatic and terrestrial. Reads were aggregated to superkingdom bins for Bacteria, Archaea, and Viruses. Within Eukaryota, reads were grouped into Fungi, Metazoa, Viridiplantae, and an aquatic algae/protists group. For Viridiplantae, reads were grouped into aquatic plants, terrestrial woody plants, and terrestrial non-woody plants, each treated as a separate conversion bin with its own trait priors.

To define the taxonomic ranges of the DNA C-value per cell and ploidy level for the stochastic model, we compiled genomic data by crawling global databases, including Animal Genome Size Database (https://www.genomesize.com/) and the Plant DNA C-values Database (https://cvalues.science.kew.org/). For each taxon, we utilized the C-value (the haploid DNA mass/cell in pg) as the primary baseline for DNA per cell. However, to reflect the actual biological state per cell, these values were adjusted for known ploidy levels. For instance, values were doubled for diploid organisms or scaled by a factor of ten or more for polyploid taxa, as provided in the C-value databases (e.g., specific aquatic algae and Viridiplantae), ensuring that the parameter represents the total DNA mass per cell rather than a theoretical haploid value.

To verify that the recovered sequencing reads are representative of total cellular DNA rather than being dominated by organellar DNA (which would bias the genome-size-based calculation), we performed a diagnostic remapping of reads against selected reference genomes for each sediment core. For each core, one interglacial and one glacial sample was selected to account for potential DNA preservation bias over time. Taxon-specific Reads from the most abundant plant and animal groups were mapped against their respective reference genomes (Supplementary Fig. 7) providing an estimate on the genomic origin of the shotgun sequenced reads. About 94–99% of reads were mapped to nuclear genomes, while contributions from organelle DNA were very small (0.03–5%). The high mapping rate justifies the use of nuclear genome size as the denominator in our organic carbon projection.

For plants and animals, globally compiled C-value datasets were available, as stated before; however, for our calculations, we selected the maximum and minimum C-values of the most abundant taxa observed in our samples (Supplementary Fig. 4, 5). For all other groups, we applied a biologically realistic range of DNA C-values (mass of DNA per cell). For carbon content estimates, a biologically realistic range of cellular carbon content values was applied to all groups (Supplementary Data 4).

Following the methodology of Herzschuh & Weiß et al.^20^, the initial DNA weight per gram of dry sediment (*DNA*_*start*_) represents the physical foundation of our calculation. This term accounts for the laboratory workflow and sediment properties:

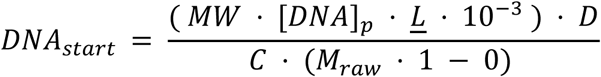

This term converts the sequencing pool concentration [*DNA*]_*p*_ back to the original sediment concentration, adjusted for molecular weight (*MW* = 325), the arithmetic mean fragment length (<u>*L*</u>), and laboratory factors (*C* & *D*). For both laboratory factors all concentration and dilution steps were recorded to determine a cumulative dilution factor (*D*) and a cumulative concentration factor (*C*).

Samples were sequenced on the Illumina NextSeq 2000 platform, achieving an average coverage of 99%. Under the assumption that the DNA on the flow cell is representatively converted into raw sequences, the taxon-specific DNA weight is estimated based on relative read percentage of the total sumcount.

In the second stage of the model, the deterministic DNA weight (*DNA*_*start*_) is multiplied by the taxon-specific read proportion (*R*_*i*_) and subjected to a Monte Carlo simulation ( *N* = 10,000 ) to estimate absolute carbon biomass. This stochastic term accounts for biological uncertainty by sampling cellular carbon content (*C*_*i*,*s*_) and DNA mass per cell (*D*_*i*,*s*_) from group-specific Beta-priors:

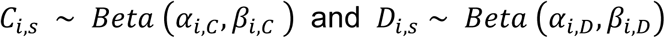

The prior database derived parameters *⍺*_*i*_ and *β*_*i*_ are tailored to the physiological characteristics of each taxon *i*. For instance, higher values of prior parameters (*⍺*_*i*,*C*_, *β*_*i*,*C*_) were assigned to woody plants compared to non-woody plants to reflect their higher carbon density.

Similarly, the distributions for prokaryotic DNA C values and cellular carbon (Supplementary Data 4) were skewed towards the lower end of the taxonomic range. This adjustment accounts for the prevalence of smaller genome sizes (DNA C-values) and lower cell-specific biomass (carbon per cell) typical of natural environments, contrasting the potentially overrepresented values in databases derived from laboratory cultures^37, 38^.

The model further incorporates an extraction efficiency factor ( *E* = 4.0 ) to correct for DNA recovery bias^36^ and structural correction factors (*σ*_*i*_) to account for non-genomic biomass components of cells or extracellular carbon (e.g., lignin or EPS (Extracellular polysaccharides))^107, 108, 109, 110, 111^. To derive a robust estimate from the stochastic process, the final biomass value (*B*^_*i*_) is defined as the *median* of all successful Monte Carlo iterations (*N* = 10,000), with the corresponding error margin represented by the interquartile range (Q1 to Q3).

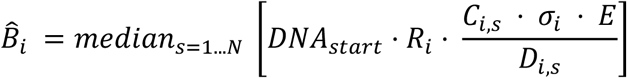

To place these biomass estimates into the context of sediment burial, burial rates (*g* ⋅ *cm*^−2^ ⋅ *yr*^−1^) were computed for each taxonomic group. Following Nürnberg et al.^112^, we multiplied the sedimentation rates (*cm* ⋅ *yr*^−1^) by the dry bulk density (*g* ⋅ *cm*^−3^). The taxon-specific biomass values (*B*^_*i*_ *wt*%) were then applied to these bulk sediment accumulation rates to derive the final carbon burial rates (OC_DNA-projected_ burial rate) for each taxonomic group.

### Model validation and sensitivity analysis

We implemented a multi-stage validation framework to ensure the robustness of the genC pipeline and to quantify the influence of individual parameters on the biomass estimates.

This included a global sensitivity analysis using the Sobol-Jansen method to identify which parameters drive the uncertainty in the final biomass estimates. We used a base sample size of, which refers to the number of variations generated for each of our four model parameters (Supplementary Fig. 14). This resulted in a total of 12,000 model evaluations. We chose this sample size to ensure that the sensitivity indices are statistically stable and that the identified drivers of uncertainty are robust. The total sensitivity index (*ST*) was calculated for four key input variables: (1) genome size (*D*_*i*,*s*_), (2) cellular carbon biomass (*C*_*i*,*s*_), (3) extraction efficiency (*E*), and (4) structural correction factors (*σ*_*i*_). Unlike local sensitivity tests, the Sobol method accounts for both the main effects and the complex interactions between these parameters. This allowed us to determine whether the variance in our biomass estimates is primarily driven by biological variability (genome size, cellular carbon biomass) or by the underlying methodological assumptions (extraction efficiency, structural correction factors).

A perturbation analysis was conducted, to evaluate the stability of the model against methodological and biological shifts. We systematically adjusted parameters in two categories. We accounted for laboratory-based and calculative uncertainties, varying extraction efficiency (*E*) and the taxon-specific structural correction factors (*σ*_*i*_) by ±10% and ±25%.

The model’s response was tested against fundamental shifts in biological trait assumptions by halving (0.5 ×) and doubling (2.0 ×) the ranges for genome size (*D*_*i*,*s*_) and cellular biomass (*C*_*i*,*s*_). This specific perturbation targets the inherent uncertainty of biological databases, which often rely on laboratory cultures that may not represent the physiological state of organisms in natural sedimentary environments. The impact of these changes in physiological states was quantified by calculating the relative change in the reconstructed median biomass (*ΔB*^_*rel*_):

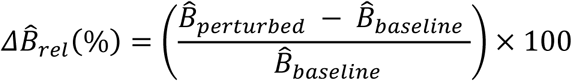

Assessing the ecological plausibility of the results was done by comparing the simulated total biomass (calculated as the sum of all taxon-specific biomass values) against measured Total Organic Carbon (TOC wt%) data. We utilized Pearson correlation (R) to evaluate trend consistency and paired Wilcoxon signed-rank tests to examine magnitude offsets. The median ratio between simulated total biomass and measured TOC served as a benchmark to determine the proportion of the total organic carbon pool represented by the reconstructed DNA-derived signal.

### Sediment TOC, and Pollen Data

Sediment total organic carbon (TOC), and pollen data for each site were obtained from previously published studies. For clarity and transparency, the source references for all datasets and specific values used in this study have been compiled in Supplementary Data 6. The TOC, as well as pollen extraction and identification, were performed following the protocols described in the cited publications.

### Statistics

We investigated the drivers of OC_DNA-projected_ burial rate and percentage of aquatic contribution to OC_DNA-projected_. We included age, average read length, sedimentation rate, vegetation compositional change, annual precipitation and July temperature as reconstructed from pollen data^113^.

To reflect the vegetation compositional change, we performed Principal Component Analysis (PCA) on the shotgun dataset of the 6 lakes including terrestrial plant families having a minimum of three values ≥ 0.01%, the lowest-abundance family included under this criterion corresponds to 1 sequencing read per family. Lake datasets were Hellinger-transformed using the decostand() function from the *vegan* package^114^ and normalized prior to PCA using the *dplyr* package^115^. Sample scores of the first axis were interpreted to follow a vegetation density gradient.

Generalized linear mixed models (GLMMs) with a Gamma error distribution and log link function were fitted using the *glmmTMB* () function from the glmmTMB package^116^ to assess the effects of environmental variables on OC_DNA-projected_ burial rate and OC_DNA-projected_ weight percentage. Model parameters were estimated using maximum likelihood, and statistical significance was assessed based on Wald z-tests. To account for site-specific variability, lake was included as a random effect in all models. All analyses were performed using R Studio (R version 4.4.2^98^). Hierarchical generalized additive models (HGAMs) were fitted using the *gam()* function from the *mgcv* package^117^ to assess temporal trends in percentage of OC_DNA-projected_ weight percentage. To highlight overall temporal patterns within the control lake, the lake was included as a random effect in the model but excluded from predictions. Statistical significance of smooth terms was evaluated using F-tests. All analyses were performed using R Studio (R version 4.4.2).

## Data and code availability

All data needed to evaluate the conclusions in the paper are present in the paper and/or the Supplementary Information. The raw shotgun sedaDNA sequence data for the six lakes originate from von Hippel et al. 2025 and Liu et al. 2025 and are deposited in the European Nucleotide Archive (ENA) (www.ebi.ac.uk/ena/browser/home) under the following bioproject numbers: Lake Ilirney (PRJEB80635) Lake Ulu (PRJEB82635), Lake Bolshoe Toko (PRJEB80642), Lake Levinson Lessing (PRJEB94536) and Lake Salmon (PRJEB82717). Lama Lake shotgun data (von Hippel et al. 2024) is deposited under PRJEB80877. The bioinformatic and statistical codes for the analyses of the data are deposited on /PolarTerrestrialEnvironmentalSystems/Yong-Weiss-et-al.-2025. (This github repository will be publicly available and converted into a Zenodo DOI after acceptance of the manuscript).

## Author contributions

U.H. conceived this study. Z.Y. and J.F.W implemented the project supervised by U.H., K.R.S. contributed to the laboratory work, conducted quality control and performed the ancient pattern analysis. S.L. performed bioinformatics. Z.Y. and J.F.W., carried out the collection of laboratory data supervised by K.R.S and U.H.. J.F.W. developed MICROSPyDER. The GenC pipeline was conceived by U.H. and J.F.W with discussion from Z.Y., K.R.S. and S.L.. J.F.W technically implemented the code of the GenC pipeline. Data analysis was performed by Z.Y., J.F.W. and S.L. supervised by U.H. The manuscript was drafted by Z.Y., J.F.W., and U.H. All authors contributed to manuscript revisions and approved the final version.

## Acknowledgements

We thank Thomas Böhmer for his help in preparing the pollen dataset. We also thank Cathy Jenks for manuscript proofreading.

