## Supplementary Materials for "Taxon-resolved sources of organic carbon preserved in lake sediments using the sedaDNA GenC pipeline"

### Content

**Supplementary Figures 1 to 24**

**Supplementary Table 1**

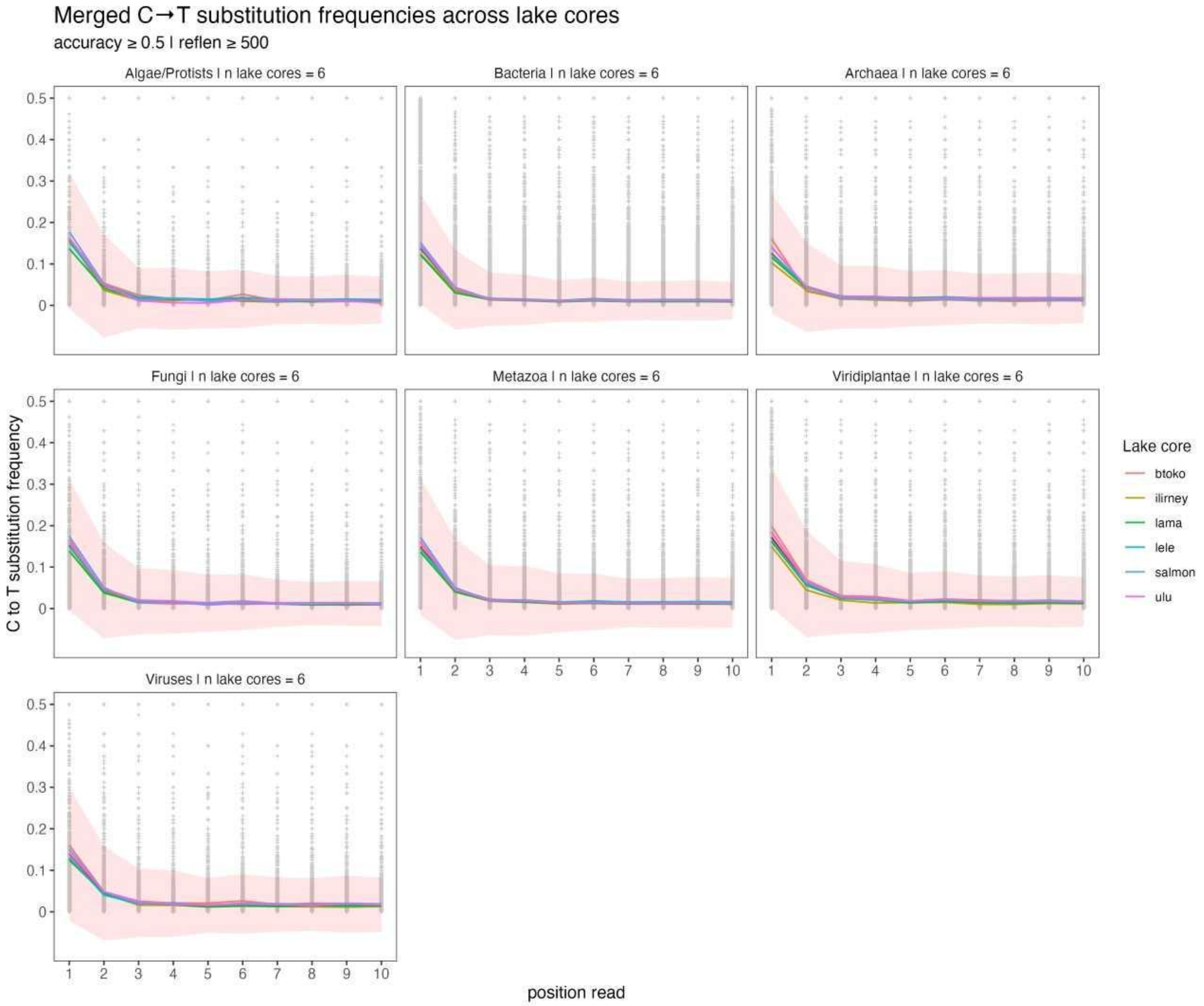


**Supplementary Figure 1. Mean C-to-T substitution frequencies for the first ten read positions of sequence reads for all taxa groups (Algae, Bacteria, Archaea, Fungi, Metazoa, Viridiplantae, Viruses) and all sites across all sample ages.** Contigs and the respective remapped sequence reads derived from the PyDamage data (per sites and taxa group) were filtered for a prediction accuracy >=0.5 and a contig length >=500 bp.

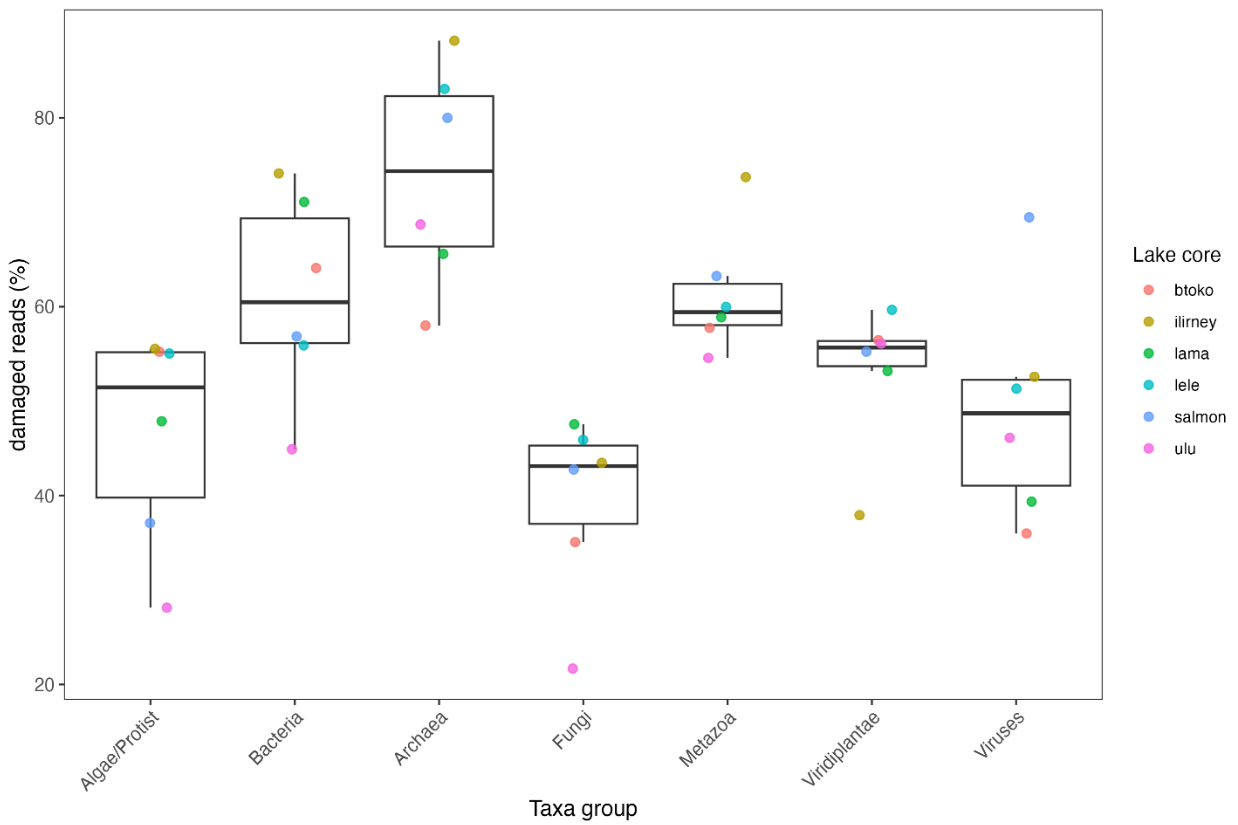


**Supplementary Figure 2. The proportion of damage (ancient) reads (from total reads) for all taxa groups and sites.** The mean % of damaged (ancient) reads is about 54.9% across all taxa groups, sites and samples ages.

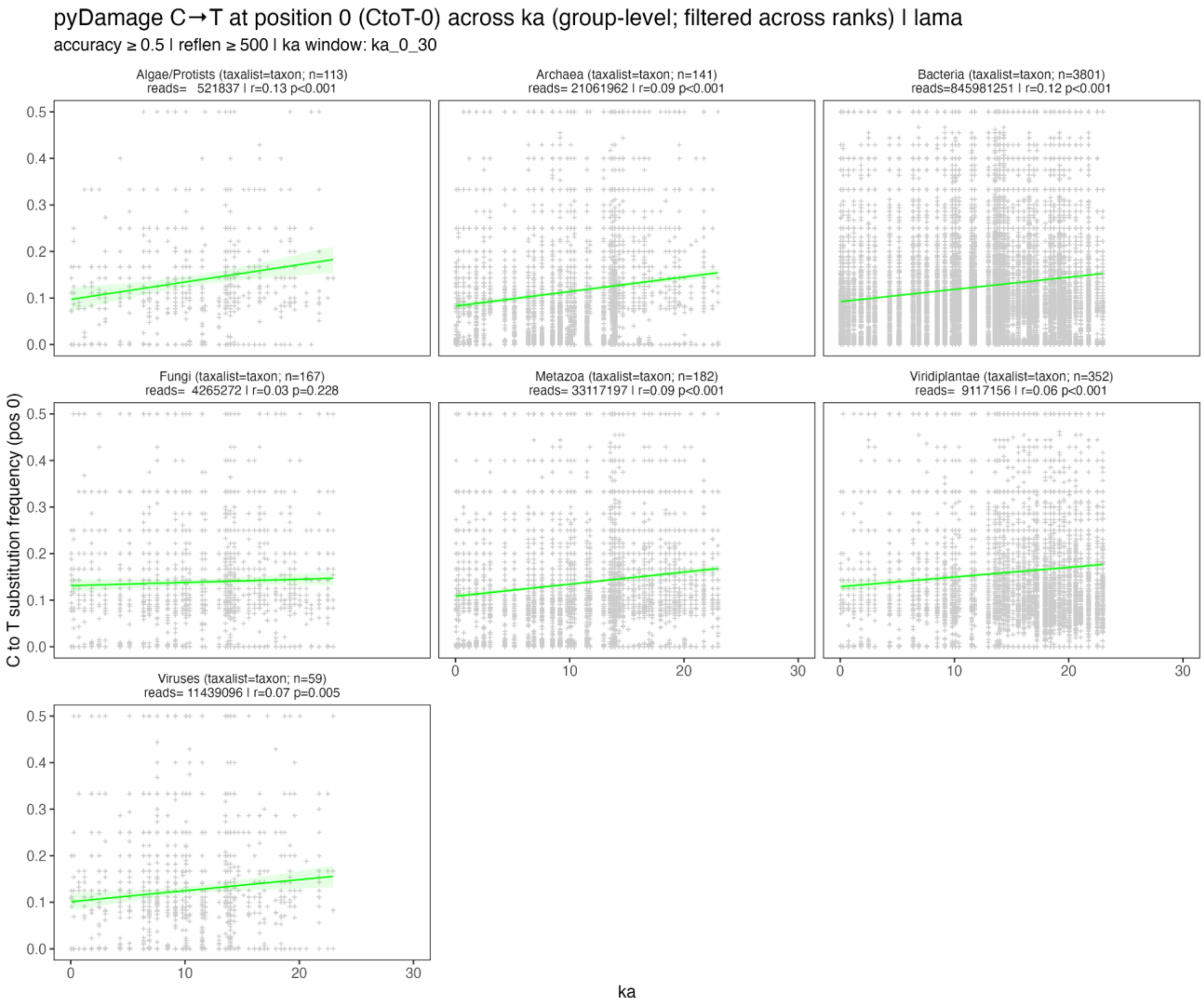


**Supplementary Figure 3. The relationship between C-to-T frequency of the first read position across age for the different taxa groups from lake Lama.**

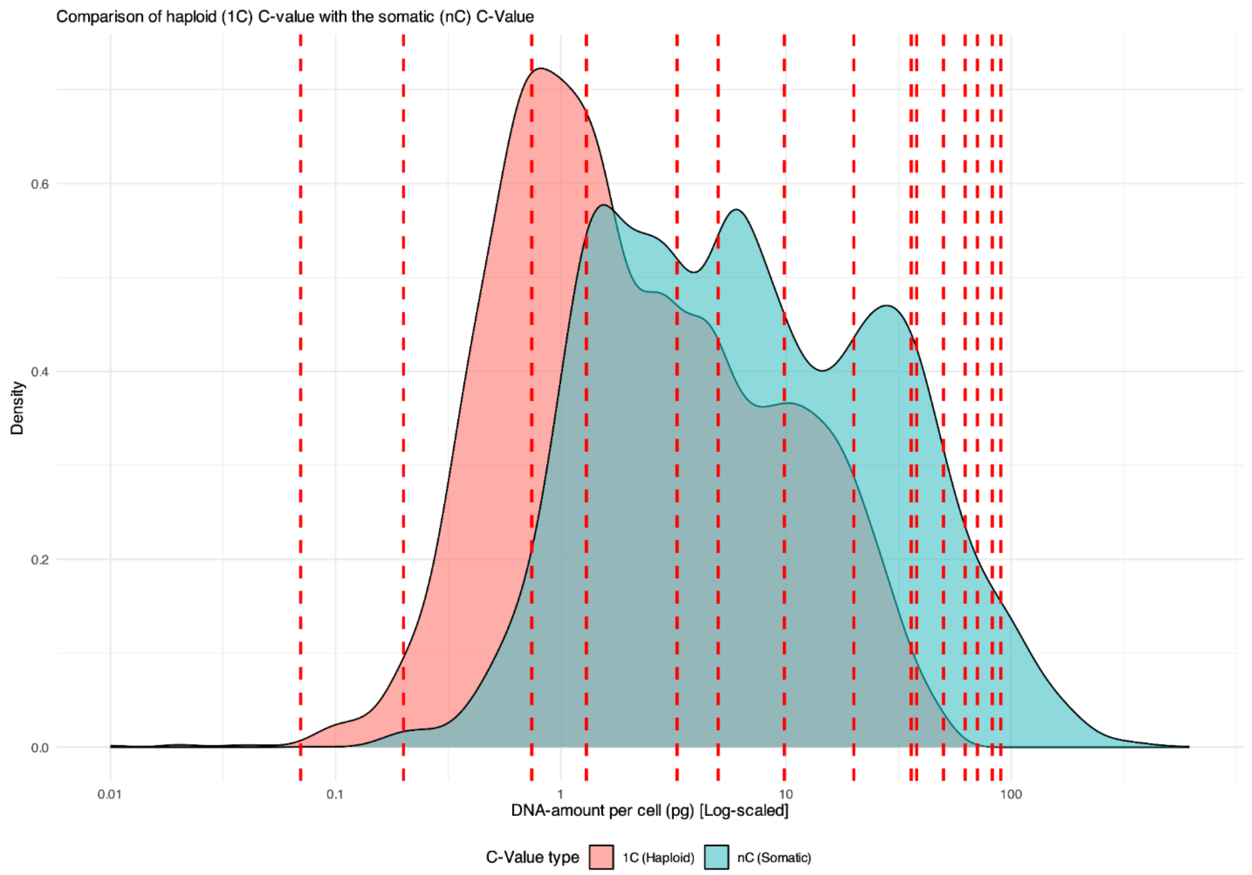


**Supplementary Figure 4. Density distributions of haploid (1C) versus somatic (nC) DNA content.** Comparison of the haploid genome size (1C, red distribution) and the estimated somatic DNA content (nC, blue distribution) across all Viridiplantae from the sedaDNA dataset, found in the database (Kew DNA C-value – https://cvalues.science.kew.org/). The x-axis displays the DNA amount per cell (pg) on a logarithmic scale to accommodate the broad spectrum of genome sizes (from 0.01 to >100 pg). The rightward shift of the density curve highlights the adjustment required to account for somatic ploidy levels in sedimentary tissues compared to baseline database values. Vertical red dashed lines indicate the data points we weighted for in the Monte Carlo Simulation.

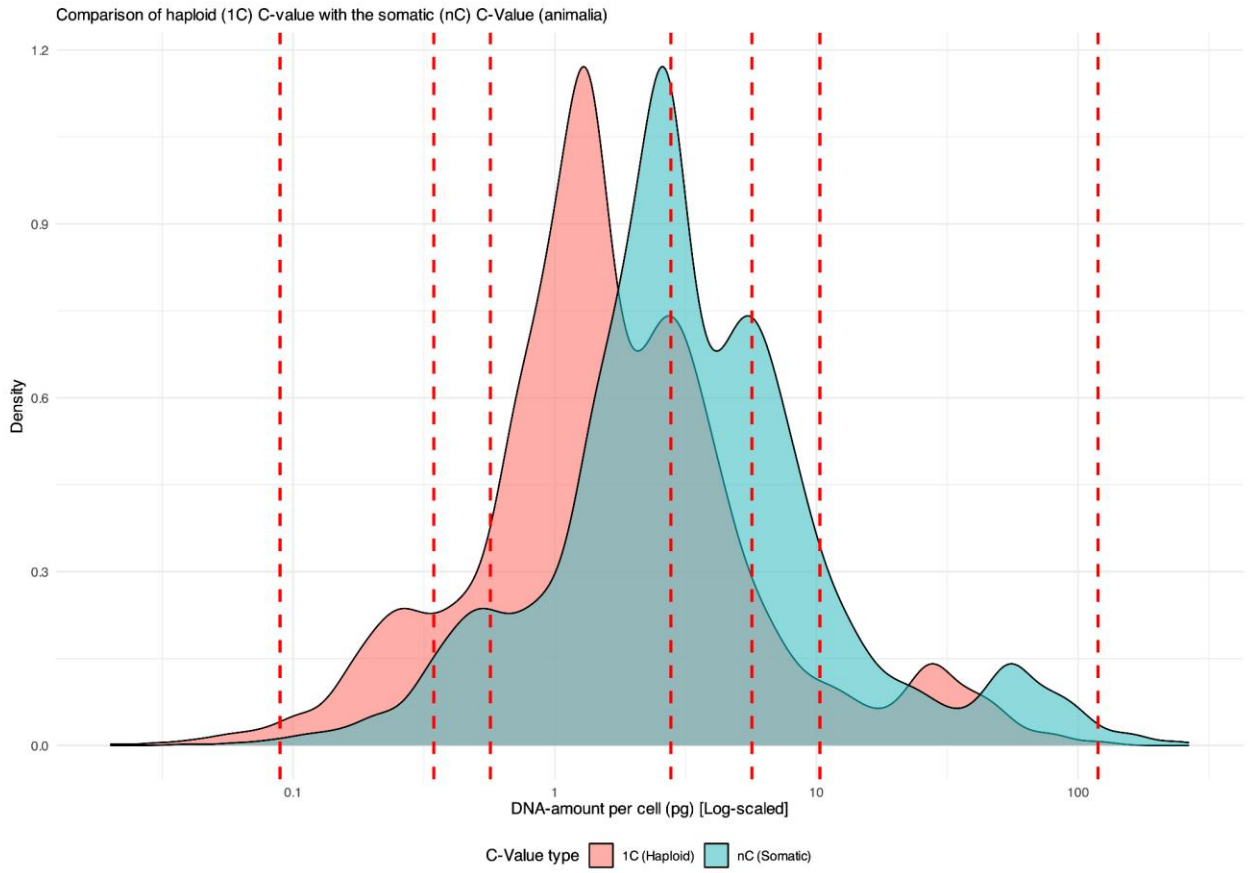


**Supplementary Figure 5. Density distributions of haploid (1C) versus somatic (nC) DNA content.** Comparison of the haploid genome size (1C, red distribution) and the estimated somatic DNA content (2C, blue distribution) across all Metazoa found in the database (Animal Genome Size – https://www.genomesize.com/). The x-axis displays the DNA amount per cell (pg) on a logarithmic scale to accommodate the broad spectrum of genome sizes (from 0.01 to >100 pg). The rightward shift of the density curve highlights the adjustment required to account for somatic ploidy levels in sedimentary tissues compared to baseline database values. Vertical red dashed lines indicate the chosen data points for the Monte Carlo Simulation.

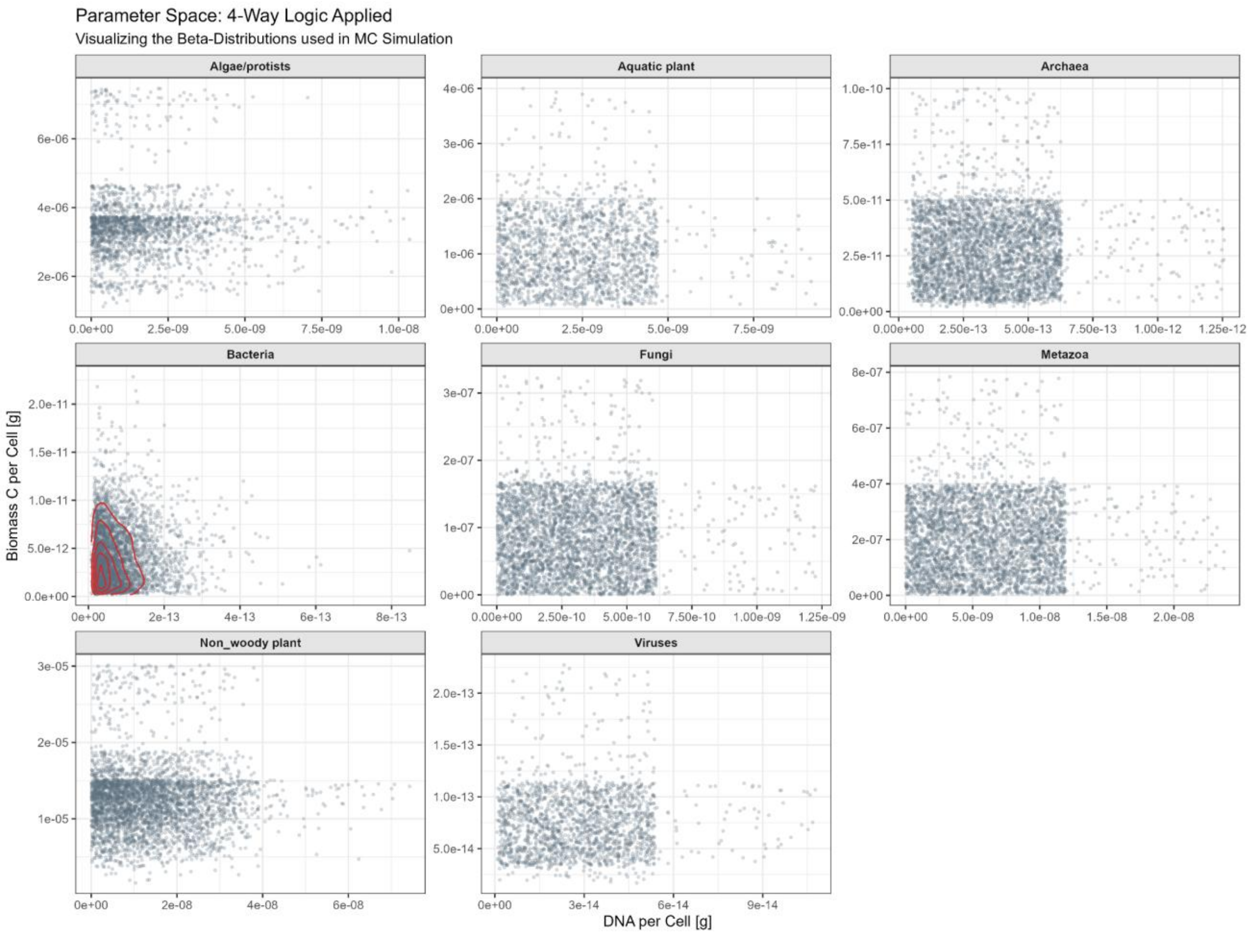


**Supplementary Figure 6.** Parameter space of DNA and biomass across organism groups under 4-way logic. This figure shows the distribution of DNA per cell (x-axis, [g]) versus biomass carbon per cell (y-axis, [g]) from Monte Carlo simulations applying the 4-way logic across different organism groups. Each point represents a simulation draw, with transparency indicating density. Panels are separated by organism groups. Red 2D density contours highlight the highest-probability regions for Bacteria. Distinct clusters reflect the scale differences among groups: Bacteria, Archaea, Viruses occupy low DNA/biomass ranges, while plants and algae show higher and more variable values, illustrating the impact of the 4-way logic on parameter distributions.

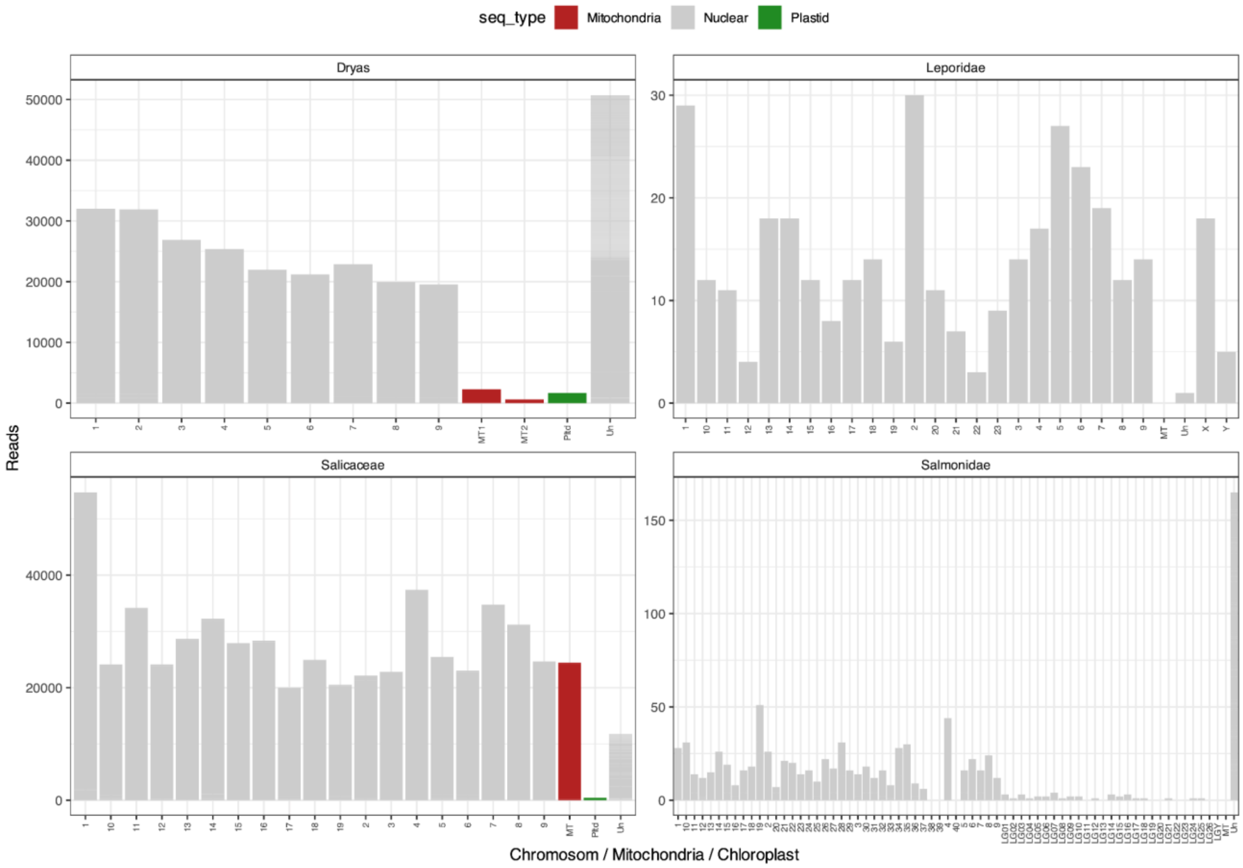


**Supplementary Figure 7. Taxon-specific remapping of dominant taxa in lake sediment cores over time.** The remapping was performed for four distinct categories: a non-woody plant (*Dryas*), a woody plant (Salicaceae), as well as limnic (Salmonidae) and terrestrial (Leporidae) Metazoa.

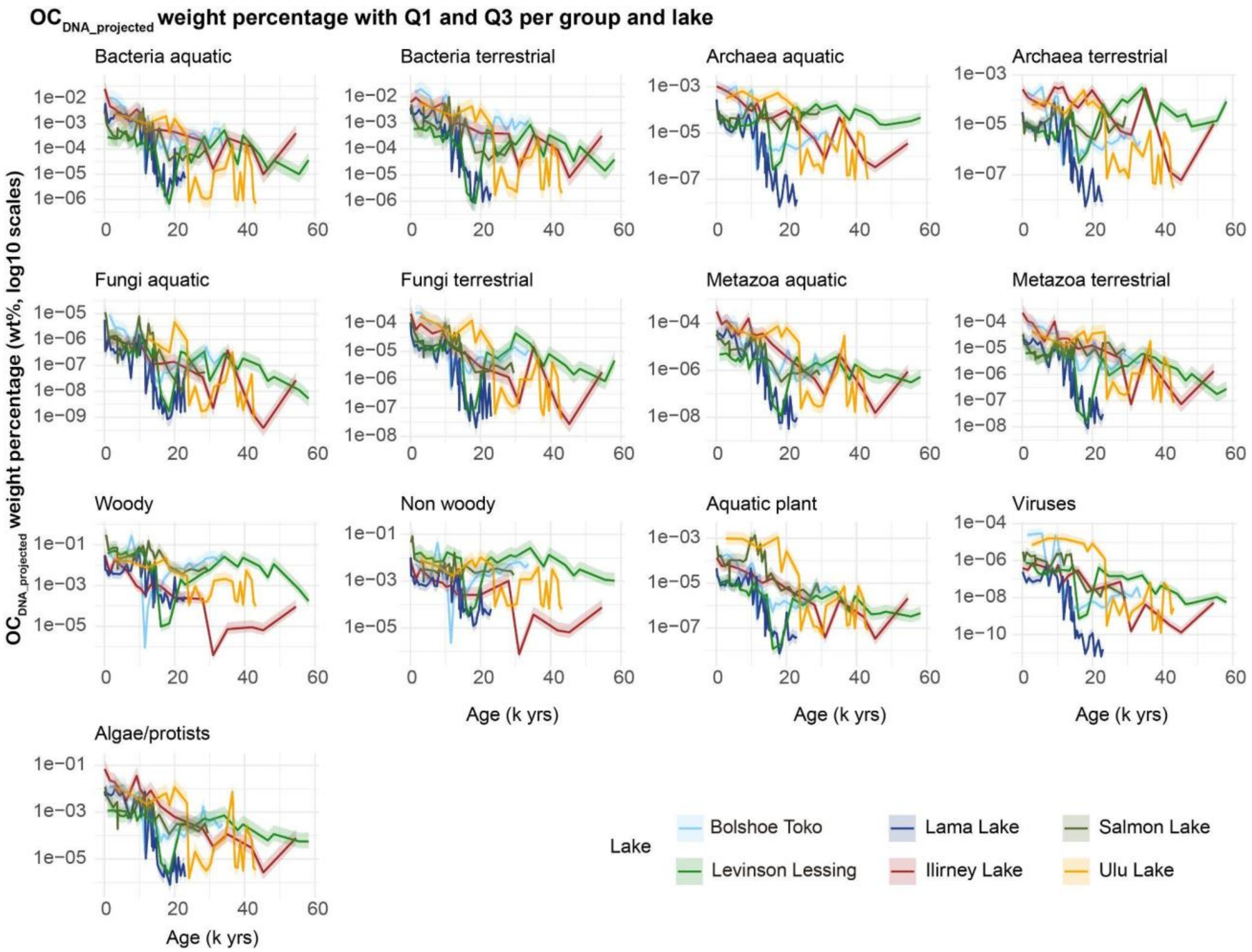


**Supplementary Figure 8. Temporal trends of OC $_{\text{DNA-projected}}$ weight percentage (wt%) across different groups and lakes.** Each panel represents a specific group, showing changes in median biomass over time. Solid lines indicate median values, while shaded ribbons represent the interquartile range (Q1–Q3), illustrating variability.

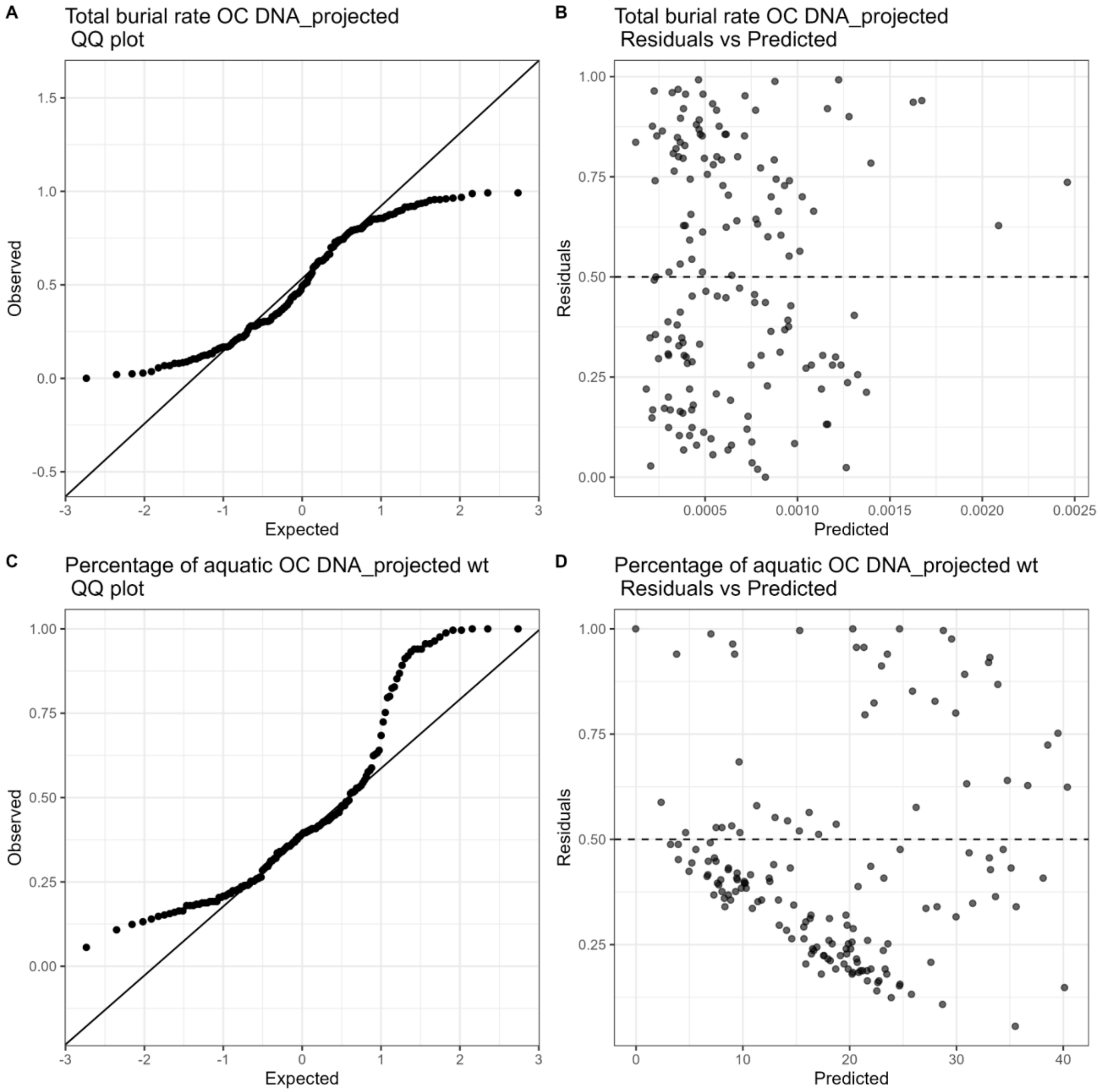


**Supplementary Figure 9. Generalized linear mixed models (GLMMs) analysis of (A–B) $OC_{DNA\ projected}$ burial rates ($g/cm^2/yr$) and (C–D) percentage of aquatic $OC_{DNA\ projected}$ weight percentage (%) and environment variables.** Panels A and C show quantile-quantile (QQ) plots comparing observed vs. expected residuals, while panels B and D show residuals versus predicted values. Both models successfully capture the main effects of environmental predictors, and residual diagnostics indicate no major deviations from model assumptions.

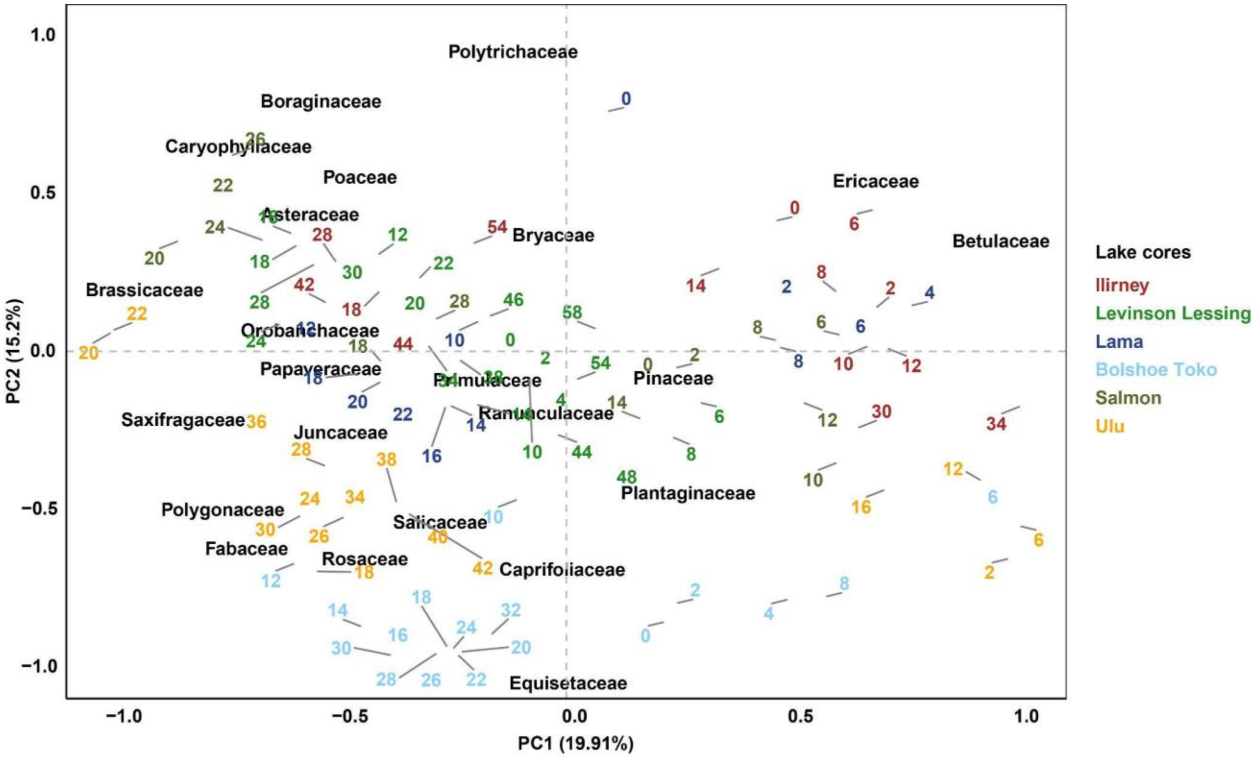


**Supplementary Figure 10. Principal Component Analysis (PCA) of DNA plant composition in the six lake cores with PC1 and PC2 scores.**

To assess patterns in terrestrial plant community composition across the different sediment cores and time intervals, we performed a Principal Component Analysis (PCA) based on relative abundances (%) of terrestrial plant taxa. The dataset included samples from 6 lake sediment cores spanning the period from 0 to 60 ka, covering major climatic intervals such as the Last Glacial Maximum, the Deglacial and the Holocene.

The data were transformed using Hellinger then scaled by lake group. The first two principal components explained 19.9123.31% and 15.217.1% of the total variance, respectively. PC1 was mainly associated with changes from Tundra to forest, while PC2 reflected variation related to humidity. Samples from the individual cores clustered separately, suggesting a distinct plant assemblage likely influenced by overall temporal and climatic shifts rather than regional climate change and local plant communities.

This analysis highlights the temporal heterogeneity of terrestrial plant community composition and provides a multivariate framework for interpreting sedaDNA-based vegetation signals across different regions.

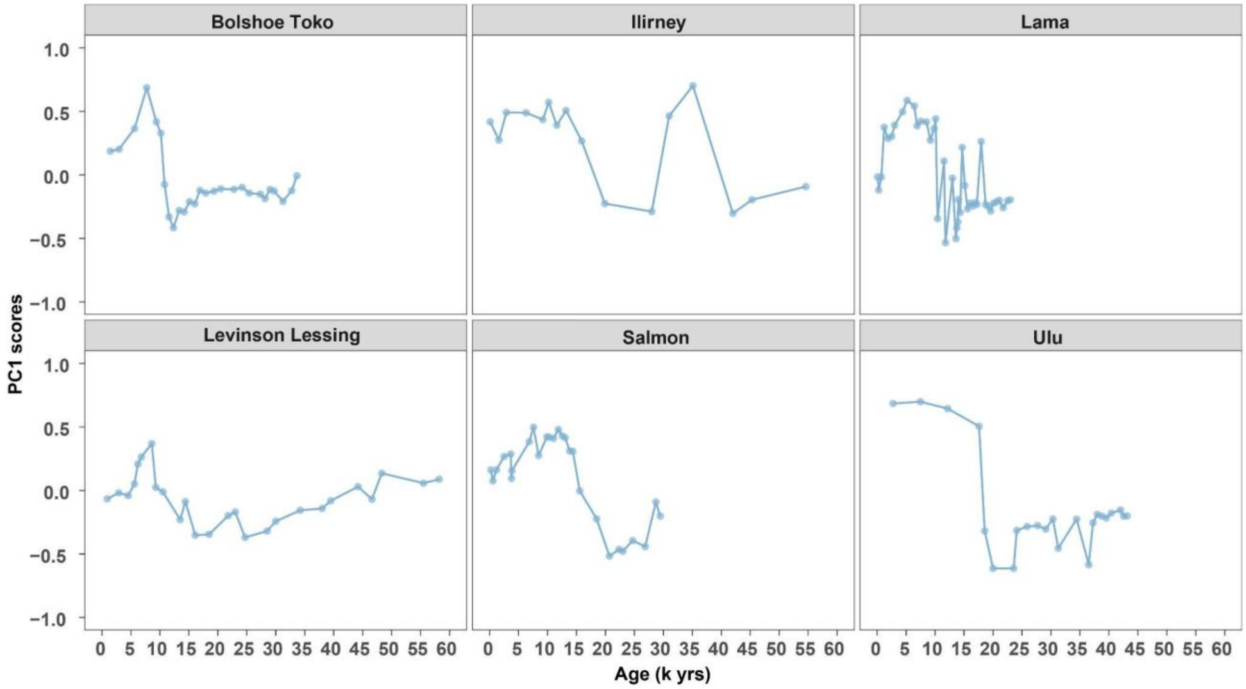


**Supplementary Figure 11. PC1 scores derived from Principal Component Analysis (PCA) of terrestrial plant composition for each lake over time.**
PC1 captures the dominant axis of variation in the dataset and reflects major shifts in vegetation composition through time. Individual cores show distinct trajectories along PC1, suggesting spatial heterogeneity in plant community responses to climatic and environmental changes. The highest lowest PC1 scores correspond to the onset of the Holocene, and may indicate transition density of vegetation.

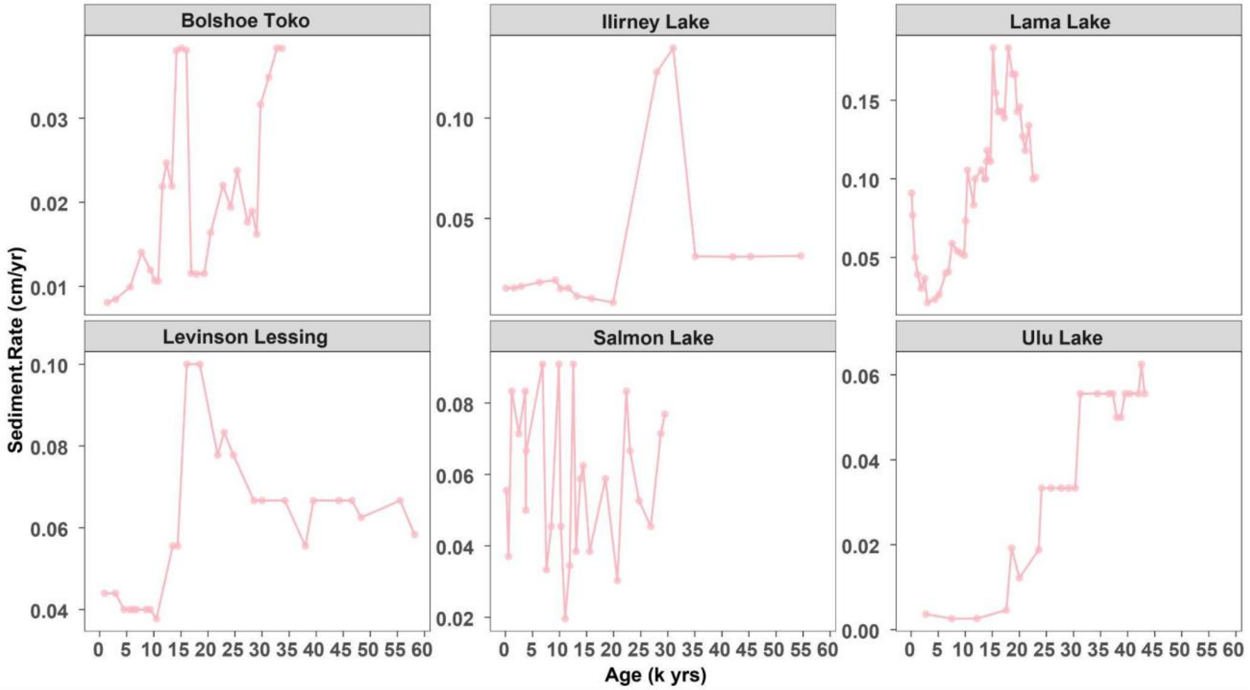


**Supplementary Figure 12. Temporal variations in sedimentation rates (cm/yr) across different lakes over time.**

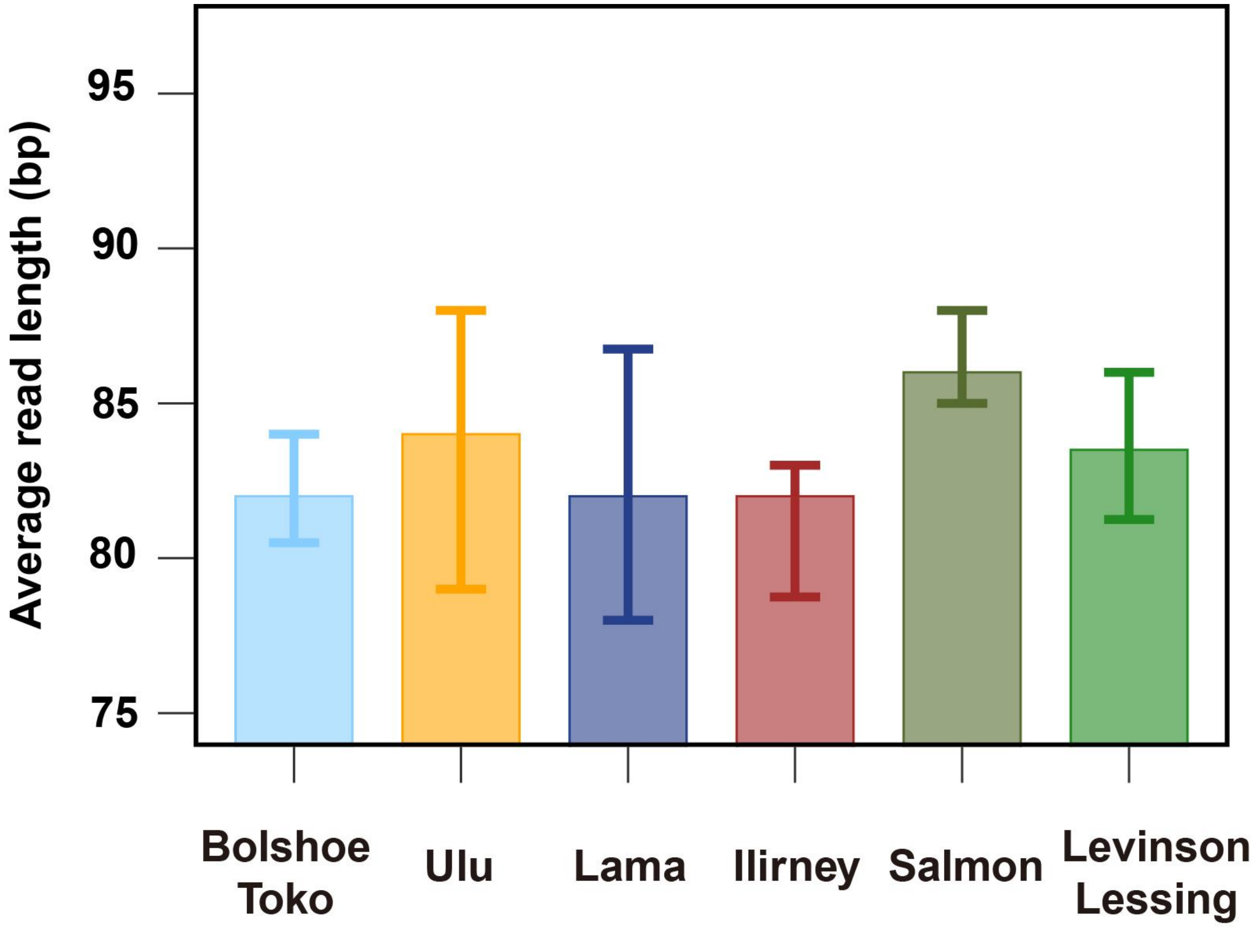


**Supplementary Figure 13. Average read length (bp) per lake.** The top of each bar represents the median of samples, while the lower and upper ends of the error bars indicate the first (Q1) and third (Q3) quartiles of those samples.

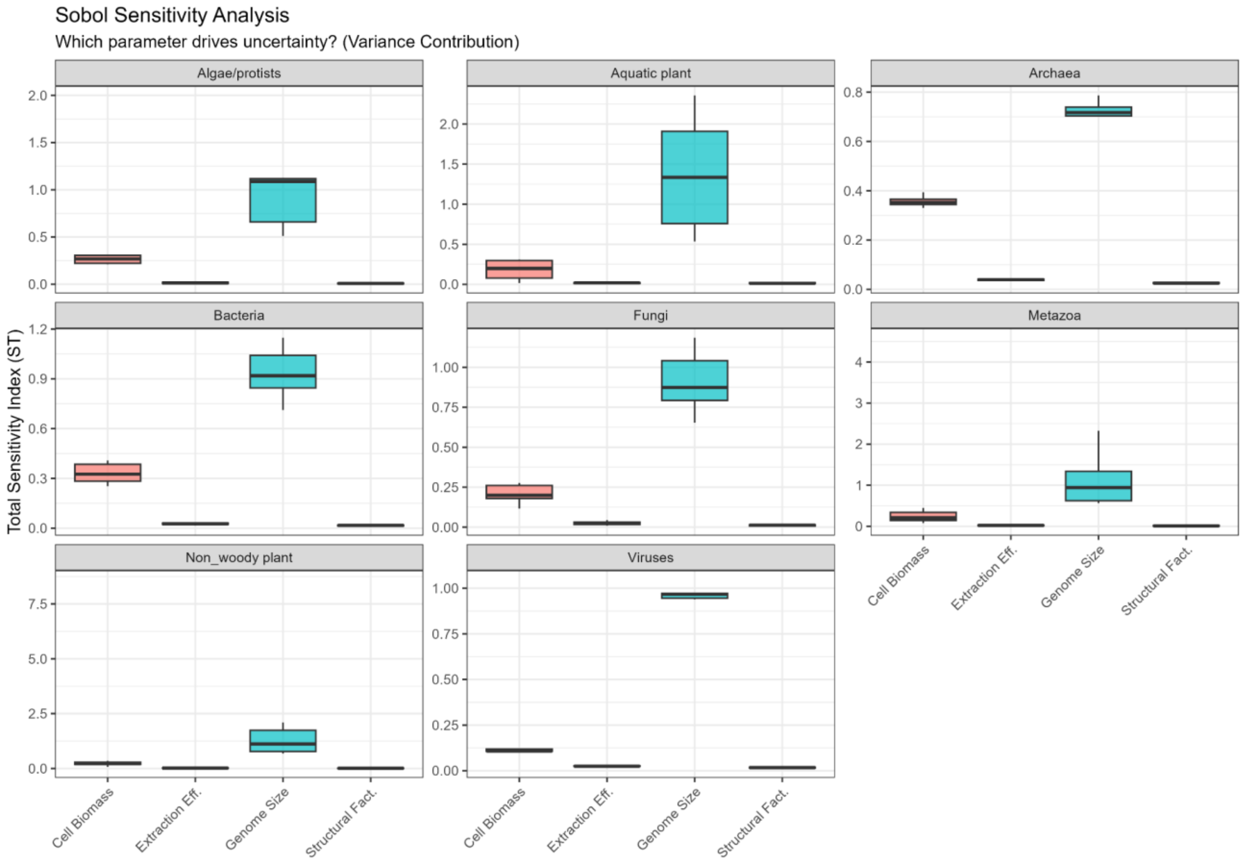


**Supplementary Figure 14. Global Sobol sensitivity analysis was conducted to evaluate the contribution of key biological and methodological parameters to the variance in $OC_{DNA\text{-}projected}$ estimates for each major taxonomic group.** Four parameters were considered for each group: cell biomass, DNA extraction efficiency, genome size, and structural factors affecting DNA recovery. Total sensitivity indices (ST) quantify the proportion of variance attributable to each parameter, including both main effects and interactions. Across all groups, genome size is the dominant contributor to variance, with secondary contributions from cell biomass. Extraction efficiency and structural factors contribute minimally. Although absolute $OC_{DNA\text{-}projected}$ values are sensitive to variation in genome size and biomass, the relative patterns among taxonomic groups remain robust.

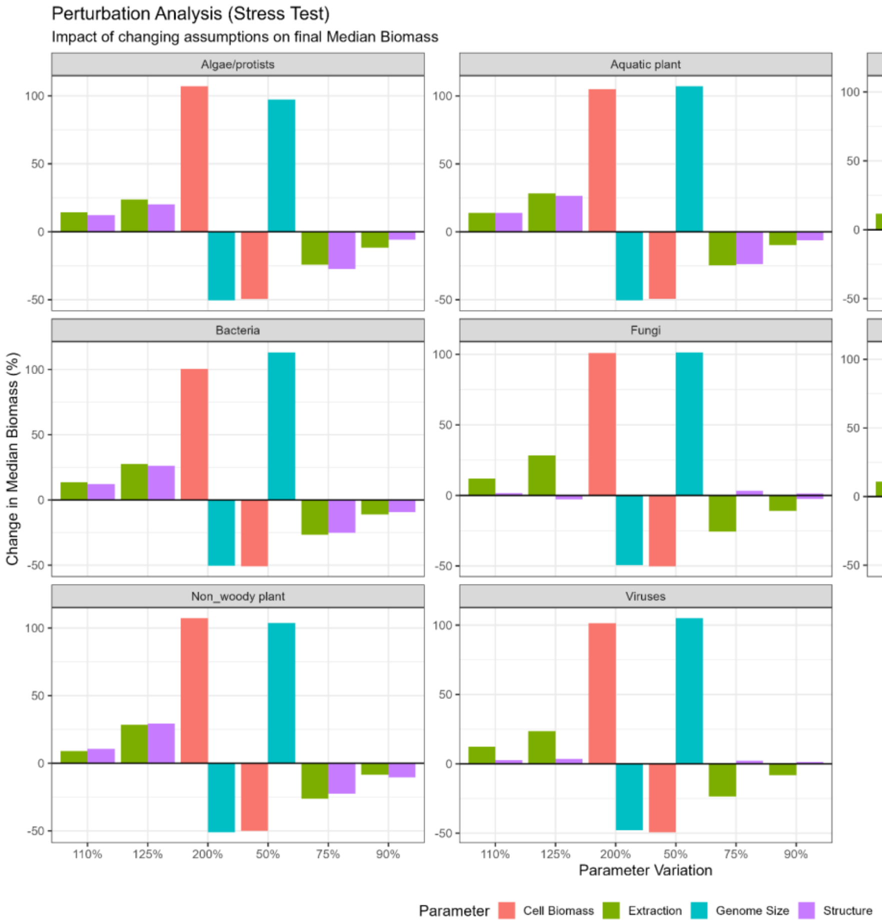


**Supplementary Figure 15. Sensitivity (perturbation) analysis of parameters used in the $OC_{DNA\text{-}projected}$ estimation pipeline.**

Parameters were independently scaled to 50%, 75%, 90%, 110%, 125%, and 200% of their baseline values, and biomass estimates were recalculated using the Monte Carlo framework. The resulting percentage change in median biomass relative to the baseline estimate (y-axis) was quantified for each taxonomic group (panels).

The analysis included both biological parameters (genome size and cell biomass) and methodological parameters (DNA extraction efficiency and structural factors). Across all taxonomic groups, biomass estimates were most sensitive to biological parameters, where a two-fold change resulted in approximately ±100% variation in estimated median biomass. In contrast, methodological parameters had substantially smaller effects, generally resulting in changes of less than ~30%.

Importantly, while parameter perturbations altered the magnitude of estimated biomass, they did not substantially change the relative ranking of the dominant taxonomic groups, indicating that the main biological conclusions are robust to variation in DNA:C assumptions.

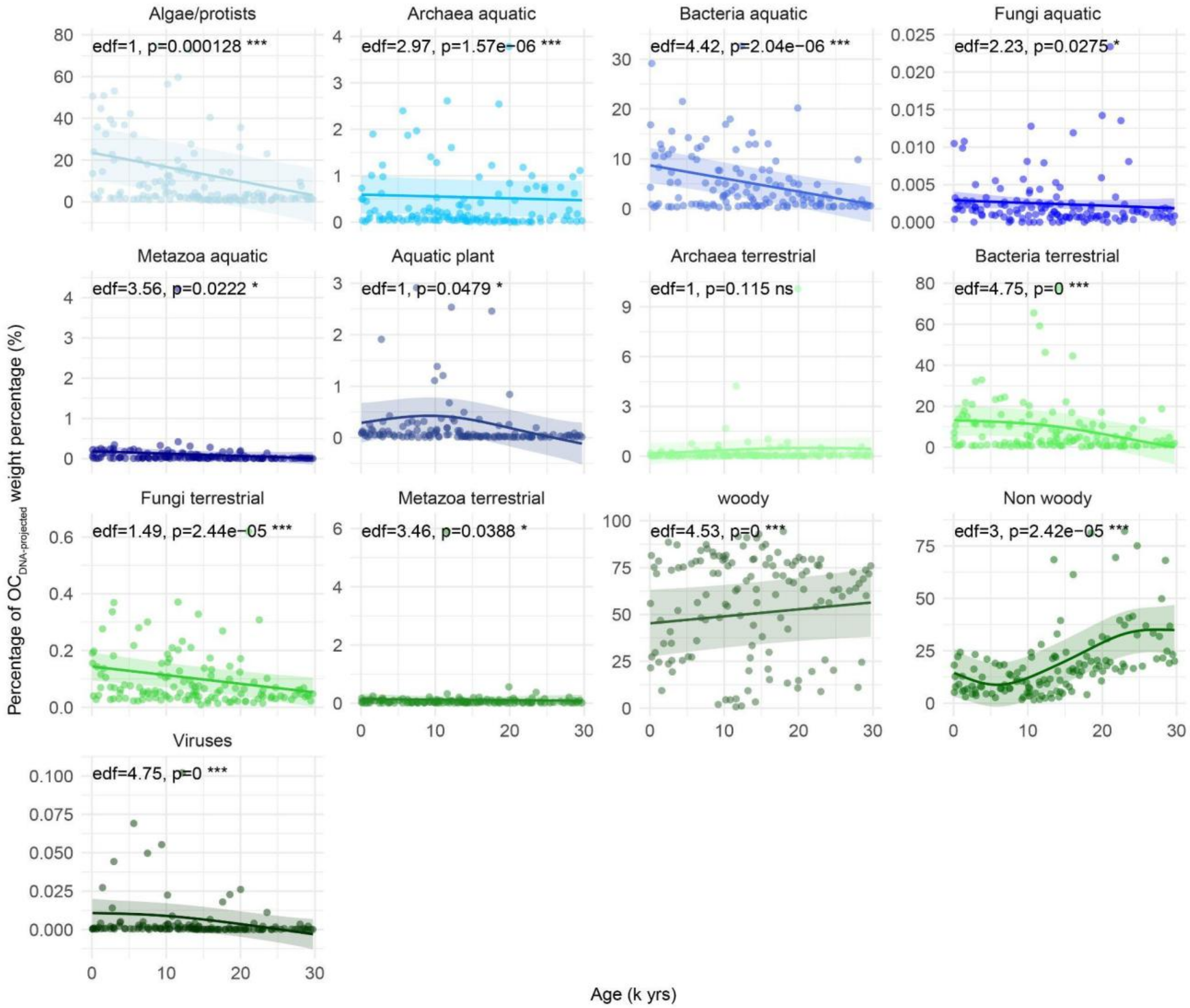


**Supplementary Figure 16. Percentage of $OC_{DNA\text{-}projected}$ weight percentage (%) for organisms from six lake cores over the past 30,000 years in the control lake.** Scatter points represent the observed Percentage of $OC_{DNA\text{-}projected}$ weight percentage (%). Solid lines show smoothed Percentage of $OC_{DNA\text{-}projected}$ weight percentage (%) trends over age fitted using Hierarchical Generalized Additive Models (HGAMs). Shaded ribbons indicate ±2 standard errors of the fitted values, approximating a 95% confidence interval. Text annotations within each panel report the estimated degrees of freedom (edf), F-statistic, and corresponding p-value for the smooth terms. The edf reflects the complexity of the fitted relationship: edf = 1 indicates a linear trend, whereas edf > 1 indicates increasing non-linearity and flexibility of the curve. Significance symbols are: *** $p < 0.001$, ** $p < 0.01$, * $p < 0.05$, and ns for not significant. Model predictions are presented with random-effect contributions excluded, in order to emphasize the underlying temporal patterns.

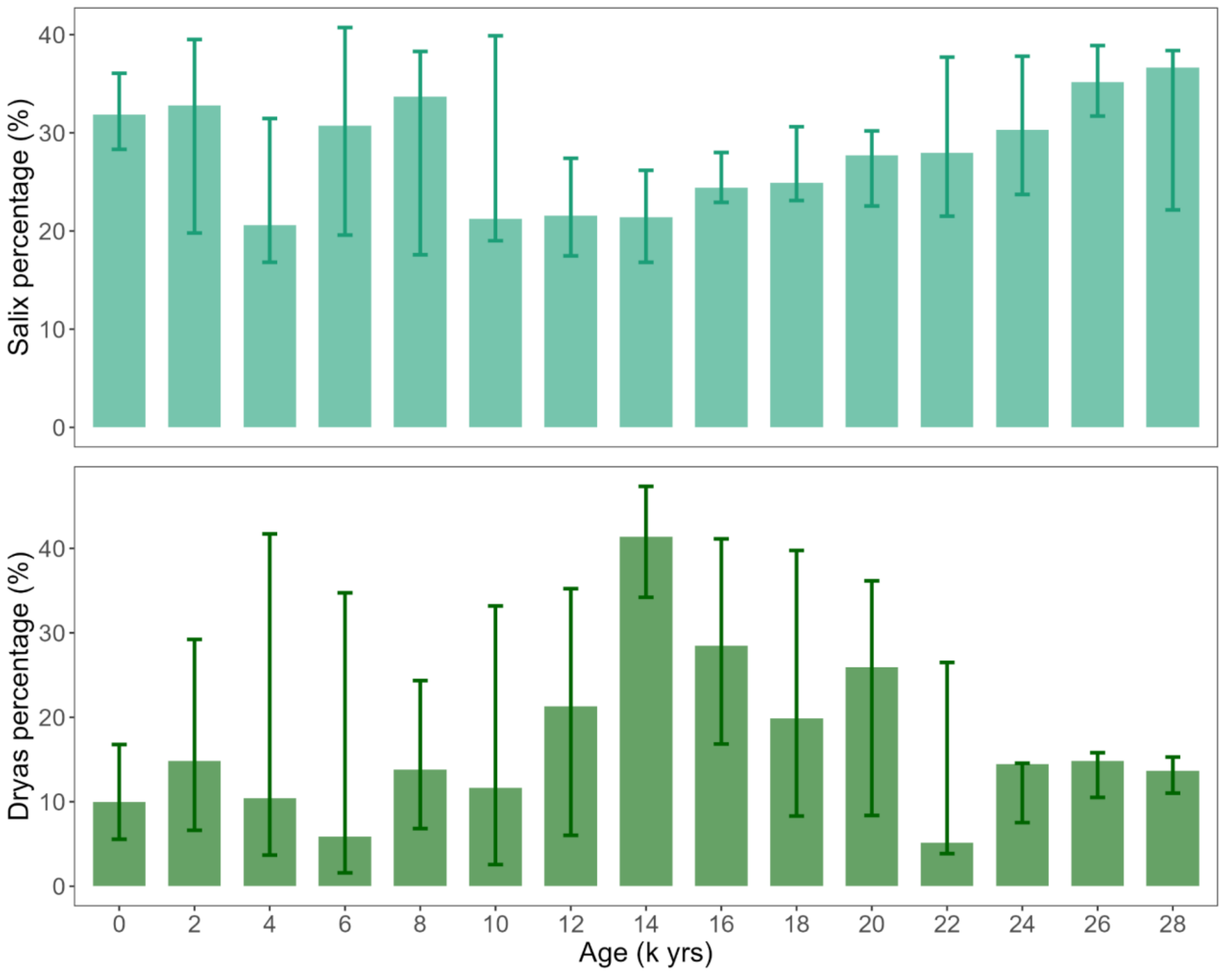


**Supplementary Figure 17. DNA reads percentages (%) of *Salix* and *Dryas* from six lake sediment cores over the past 30,000 years.** The top of each bar represents the median of samples within each 2000-year interval, while the lower and upper ends of the error bars indicate the first (Q1) and third (Q3) quartiles of those samples. Bars are plotted at the beginning of each interval.

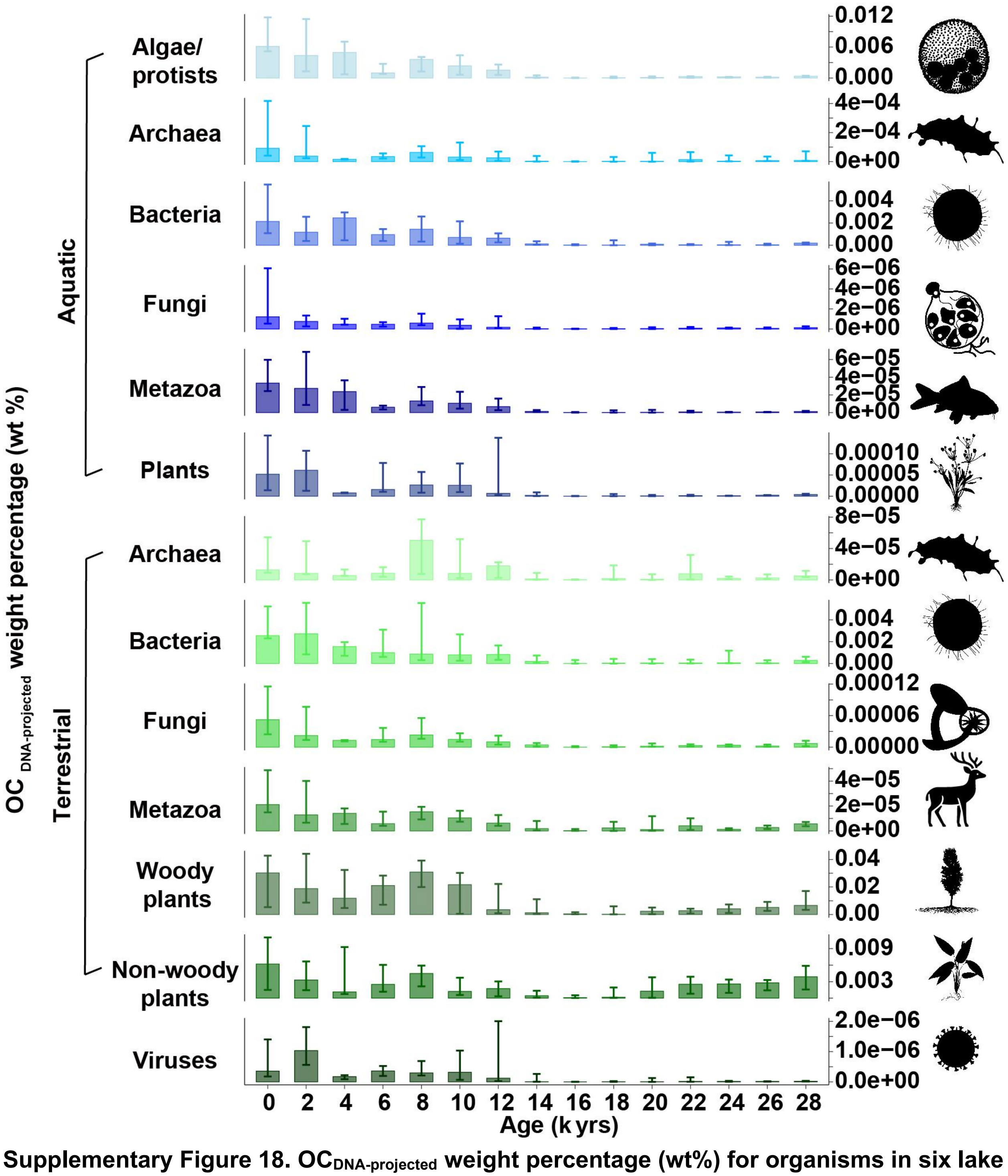


**Supplementary Figure 18. $OC_{DNA\text{-}projected}$ weight percentage (wt%) for organisms in six lake cores over the past 30,000 years.** The top of each bar represents the median of samples within each 2000-year interval, while the lower and upper ends of the error bars indicate the first (Q1) and third (Q3) quartiles of those samples. Bars are plotted at the beginning of each interval.

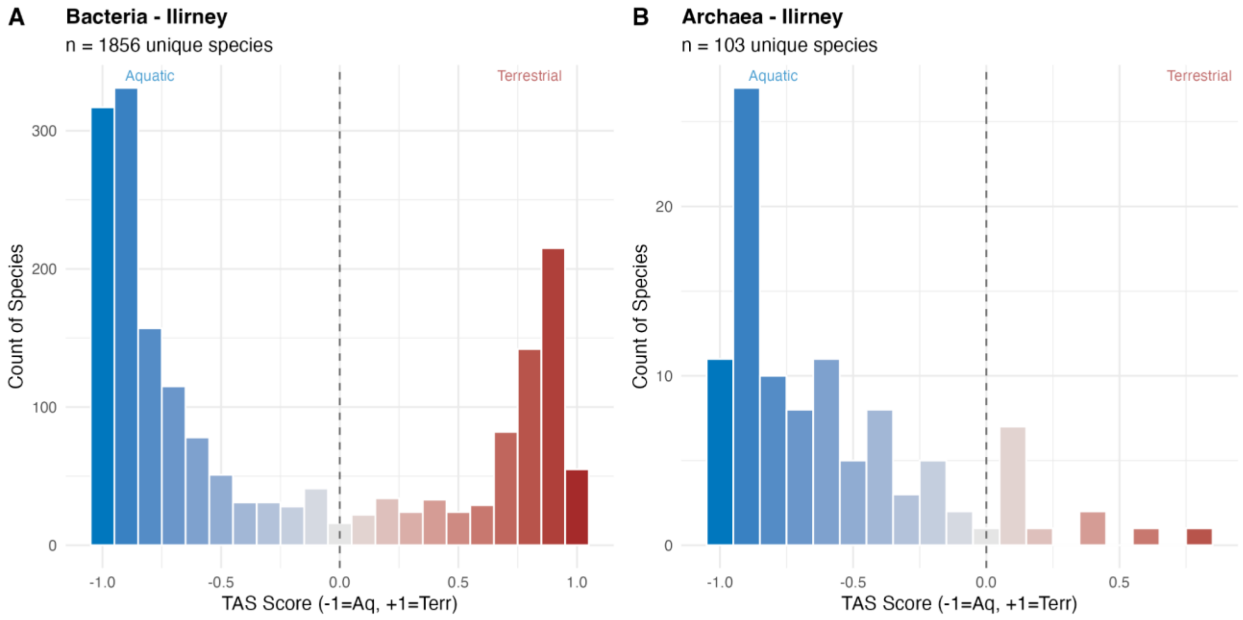


**Supplementary Figure 19. Distribution of Habitat Affinity Scores (HAS) for prokaryotic communities in the Ilirney sediment core.** The histograms show the classification of **(A)** Bacteria (n=1856 unique species) and **(B)** Archaea (n=103 unique species) based on their habitat preference. The HAS ranges from −1.0 (strictly aquatic, blue) to +1.0 (strictly terrestrial, red). The dashed vertical line at 0.0 indicates the threshold between predominantly aquatic and terrestrial taxa.

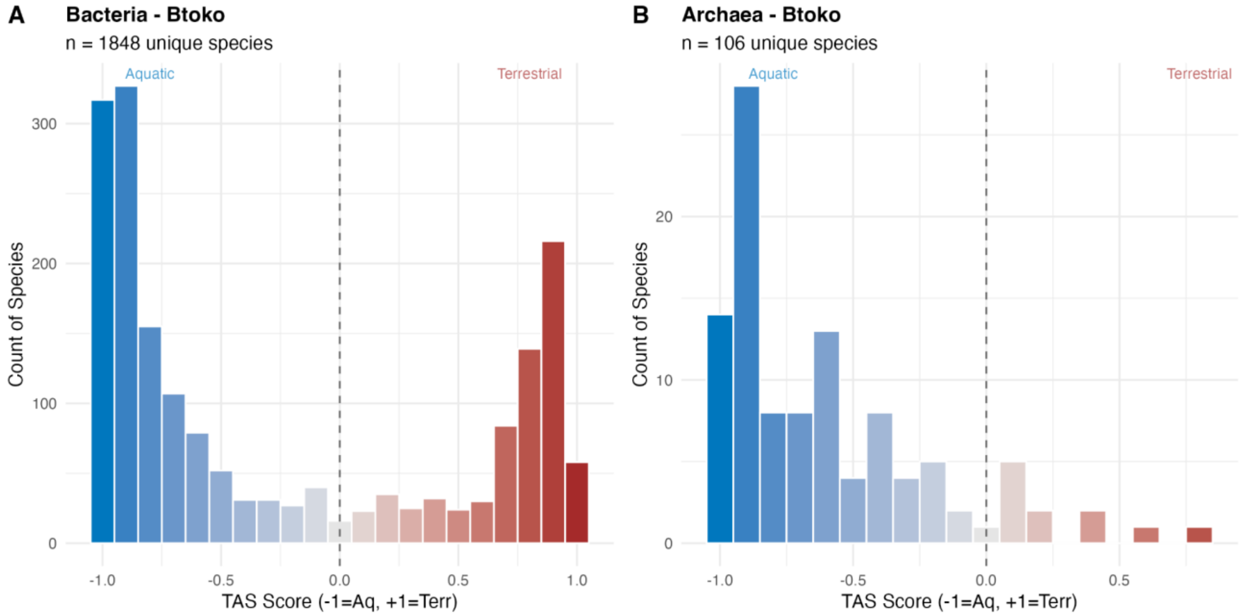


**Supplementary Figure 20. Distribution of Habitat Affinity Scores (HAS) for prokaryotic communities in the Bolshoe Toko (BToko) sediment core.** The histograms show the classification of **(A)** Bacteria (n=1848 unique species) and **(B)** Archaea (n=106 unique species) based on their habitat preference. The HAS ranges from −1.0 (strictly aquatic, blue) to +1.0 (strictly terrestrial, red). The dashed vertical line at 0.0 indicates the threshold between predominantly aquatic and terrestrial taxa.

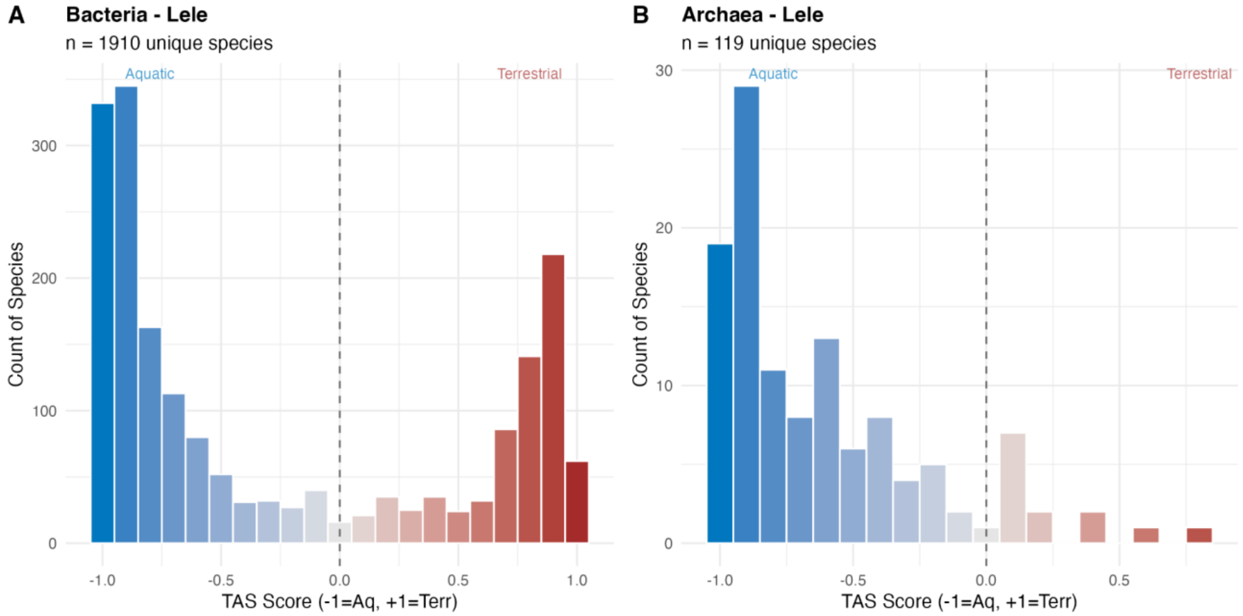


**Supplementary Figure 21. Distribution of Habitat Affinity Scores (HAS) for prokaryotic communities in the Levinson Lessing (Lele) sediment core.** The histograms show the classification of **(A)** Bacteria (n=1910 unique species) and **(B)** Archaea (n=119 unique species) based on their habitat preference. The HAS ranges from −1.0 (strictly aquatic, blue) to +1.0 (strictly terrestrial, red). The dashed vertical line at 0.0 indicates the threshold between predominantly aquatic and terrestrial taxa.

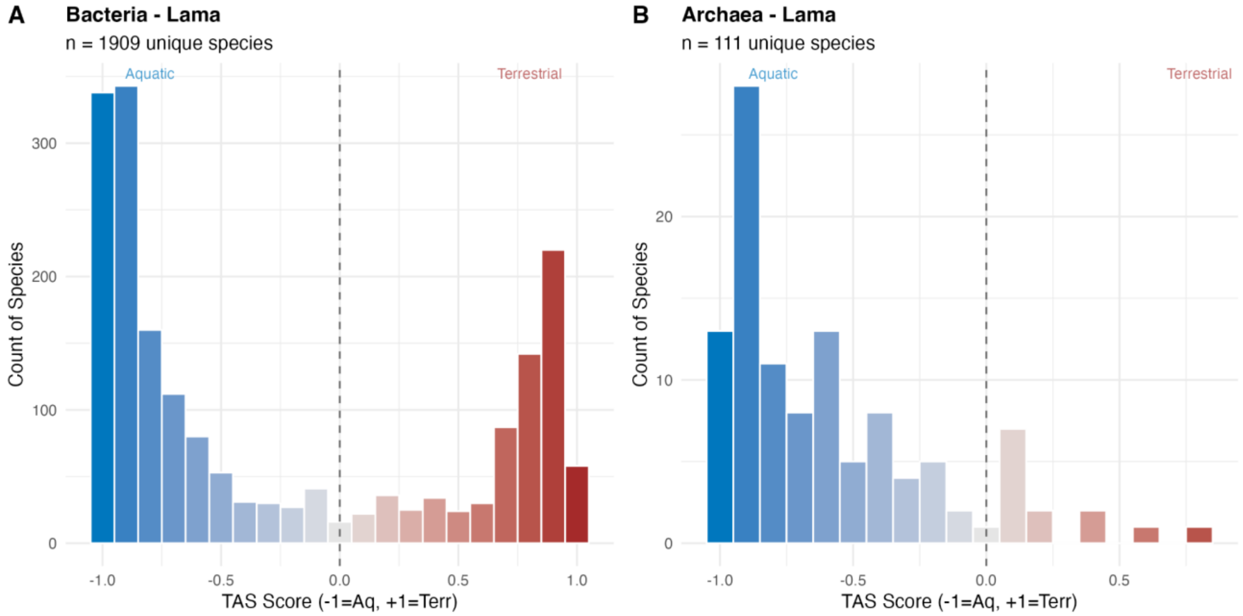


**Supplementary Figure 22. Distribution of Habitat Affinity Scores (HAS) for prokaryotic communities in the Lama sediment core.** The histograms show the classification of **(A)** Bacteria (n=1909 unique species) and **(B)** Archaea (n=111 unique species) based on their habitat preference. The HAS ranges from −1.0 (strictly aquatic, blue) to +1.0 (strictly terrestrial, red). The dashed vertical line at 0.0 indicates the threshold between predominantly aquatic and terrestrial taxa.

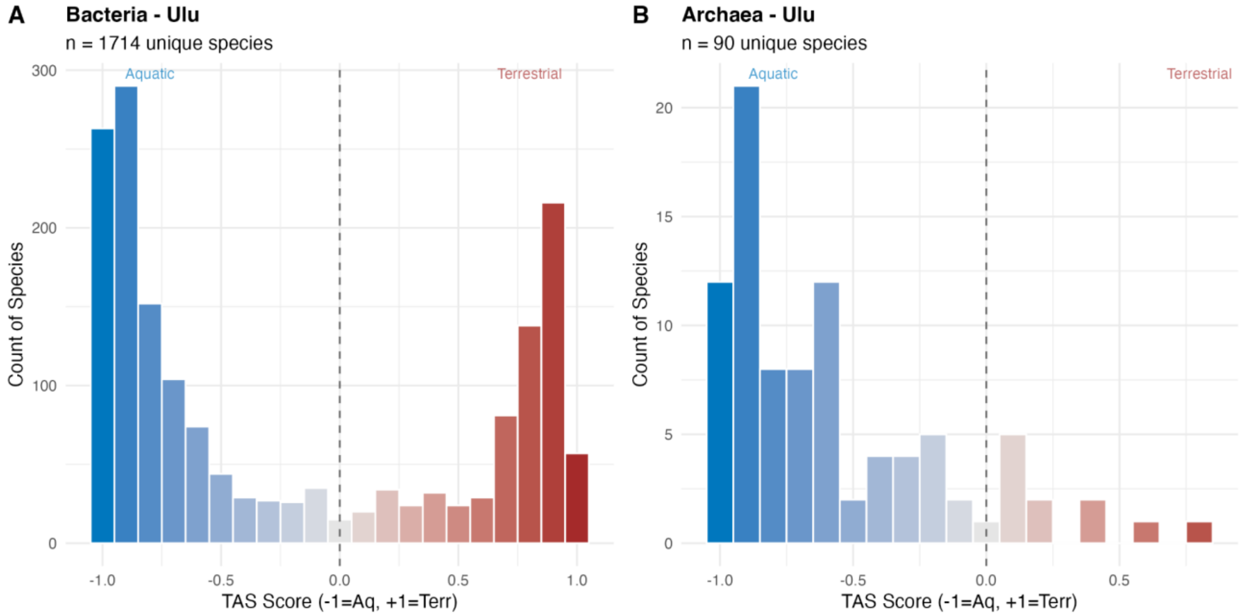


**Supplementary Figure 23. Distribution of Habitat Affinity Scores (HAS) for prokaryotic communities in the Ulu sediment core.** The histograms show the classification of **(A)** Bacteria (n=1714 unique species) and **(B)** Archaea (n=90 unique species) based on their habitat preference. The HAS ranges from −1.0 (strictly aquatic, blue) to +1.0 (strictly terrestrial, red). The dashed vertical line at 0.0 indicates the threshold between predominantly aquatic and terrestrial taxa.

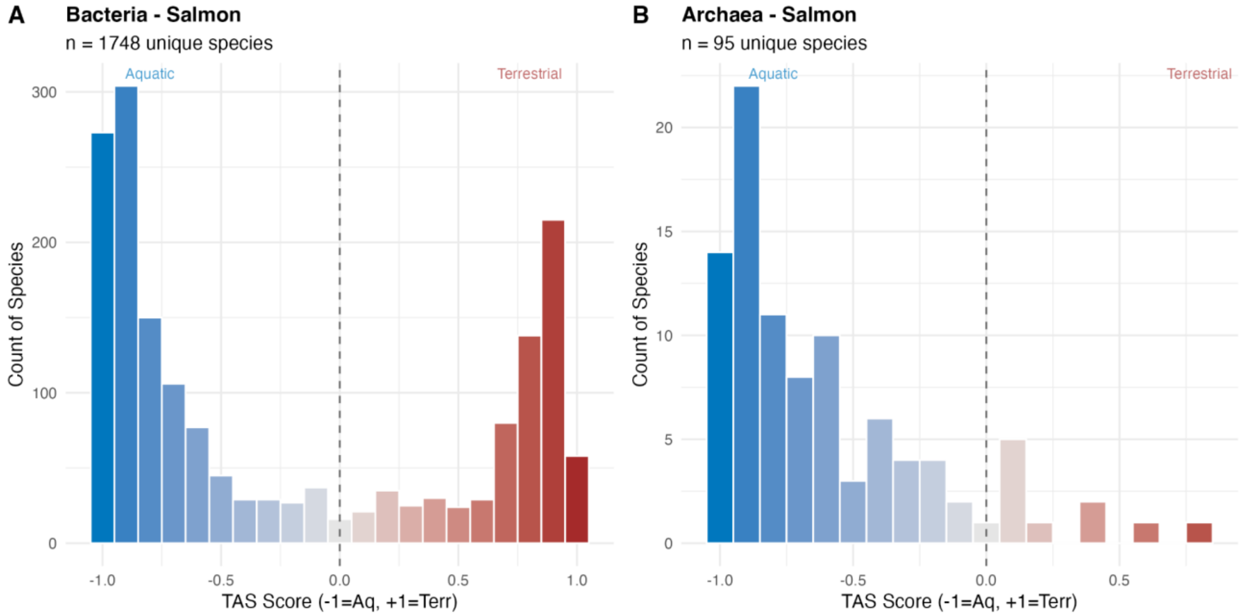


**Supplementary Figure 24. Distribution of Habitat Affinity Scores (HAS) for prokaryotic communities in the Salmon sediment core.** The histograms show the classification of **(A)** Bacteria (n=1748 unique species) and **(B)** Archaea (n=95 unique species) based on their habitat preference. The HAS ranges from −1.0 (strictly aquatic, blue) to +1.0 (strictly terrestrial, red). The dashed vertical line at 0.0 indicates the threshold between predominantly aquatic and terrestrial taxa.

**Supplementary table 1.** Main characteristics of the sampled lakes.

| Lake | Coordinates | pH | Electrical conductivity (µS $cm^{-1}$) | Dimension | Reference |
|---|---|---|---|---|---|
| Bolshoe Toko | 56.15 °N, 130.30° E; 903 m a.s.l | 6.8 | 35.1 | area: 15.4 km*7.5 $km^2$, maximum depth: 72.5 m | (Biskaborn et al., 2021)[1] |
| Ulu | 63.35 °N, 141.08° E; 950 m a.s.l | 7.6 | 73.4 | area: 4.8 $km^2$, maximum depth: approximately 35–40 m | (Jia et al., 2024)[2] |
| Lama | 69.32 °N, 90.12 °E; 53 m a.s.l. | 7.55 | 96 | area: 466 $km^2$; maximum depth: 254 m (Andreev, 2004) | (Hahne and Melles, 2006)[3] |
| Ilirney | 67.35 °N, 168.3167°E; 407 m.a.s.l. | 8.43 | 62 | area: 29.7 $km^2$; maximum depth: 44 | (Vyse et al., 2020)[4] |
| Salmon | 64.91 °N, 164.989 °E; 135 m.a.s.l. | 7.2 | 220 | area: 1.1 $km^2$ maximum depth: 21 | Submission to Pangaea ongoing |
| Levinson-Lessing | 74.27 °N, 98.39 °E; 48 m a.s.l | 7.6 | 120 | 15 km * 2.5 km, maximum depth: 120 m | (Lebas et al., 2019; Melles et al. 1996,)[5,6] |
